# Target-driven optimization of feature representation and model selection for microbiome sequencing data with *ritme*

**DOI:** 10.64898/2025.12.08.693045

**Authors:** Anja Adamov, Christian L. Müller, Nicholas A. Bokulich

## Abstract

Microbiome sequencing datasets are sparse, high-dimensional, compositional, and hierarchically structured, and predictive modeling from them typically relies on *ad hoc* feature representation choices that obscure their impact on performance and interpretation. We present *ritme*, an open-source Python package that jointly optimizes microbiome-specific feature representation and model selection — combined algorithm selection and hyperparameter optimization — tailored to these data. *ritme* systematically searches taxonomic aggregation, sparsity-aware selection, compositional transforms, and metadata enrichment together with model class and hyperparameters, using state-of-the-art optimizers that scale from a laptop to a compute cluster. Across three real-world use cases, *ritme* outperformed the original study pipelines by 7–29% on the primary task metric and surpassed three AutoML baselines in six of seven comparisons, while selecting substantially fewer features and exposing how feature and model choices drive performance. Open-source and modular, *ritme* supports reproducible, parsimonious predictive modeling, downstream biological investigation, and extension to other multi-omics modalities.

**Highlights:**

- Open-source *ritme* co-optimizes microbiome feature representation and model selection
- Outperforms expert-designed pipelines by 7–29% and AutoML baselines in 6 of 7 cases
- Selects far fewer features and quantifies feature and model contributions
- Scales across compute clusters and extends to other omics modalities and model types

**In brief.:** Predictive modeling from microbiome sequencing data relies on *ad hoc* feature choices that obscure performance and reproducibility. *ritme* is the first open-source framework to jointly optimize microbiome-specific feature engineering — taxonomic aggregation, sparsity-aware selection, compositional transforms, and metadata enrichment — with model selection and hyperparameters, using optimizers that scale to distributed compute clusters. Across three real-world studies, *ritme* outperforms expert-designed pipelines and AutoML baselines in six of seven comparisons, while exposing how feature and model choices drive performance.

## Introduction

Microbiomes are complex ecosystems of microorganisms and their functional components, which play key roles in human health, agriculture, and the environment. In human health, microbial dysbiosis has been implicated in inflammatory bowel disease^1^ and in numerous immunological and metabolic conditions^2,3^. In agriculture, microbiome composition governs soil health^4^, wine quality^5^, and crop performance^6^. In environmental systems, microbiomes play important roles in food chains as well as biogeochemical cycling^7^, and shifts in microbial compositions are associated with temperature^8^ and salinity gradients^9^. Discovering new functional roles and interactions within complex microbiomes can be challenging due to the high dimensionality as well as the large proportion of uncharacterized species and functions present in many ecosystems. Hence, identifying microbial signatures associated with these covariates typically involves training machine learning (ML) models on microbial features to predict selected targets such as disease states, environmental parameters or agronomic traits^10,11^. In these models, effective feature representation and model choice are crucial for obtaining robust predictive performance^12–15^, particularly given the limited sample sizes relative to the large number of features typical of microbiome datasets^16^. The resulting predictive models then serve as a basis for further investigation of the underlying biology.

Standardized bioinformatics pipelines exist for processing raw microbial nucleotide sequence reads into feature count tables of similar sequence clusters^17–20^ or error-corrected reads^21^. However, there is a lack of user-friendly and optimized workflows for their downstream processing that address all statistical properties inherent to microbial count data — including sparsity, high dimensionality, compositionality^22^, and phylogenetic and functional hierarchies — while optimizing predictive performance. Researchers often make subjective, non-reproducible decisions about microbial feature aggregation, selection, and transformation prior to model optimization, obscuring how these influence predictive performance and the biological insights obtained^16,23^. Alternatively, they address only selected statistical properties with microbiome-specific feature representation methods restricted to fixed, manually defined cutoff values evaluated via grid-search^15,24^ or evaluated on a predefined predictive model algorithm and associated hyperparameters^25^.

Many automated ML (AutoML) pipelines^26–29^ perform Combined Algorithm Selection and Hyper-Parameter Optimization (CASH) to optimally find the best model algorithm and feature preprocessing steps. Their feature processing repertoire is largely confined to generic statistical operations (e.g., constant variance feature removal and standardization^28^) and dimensionality reduction techniques (e.g., feature embeddings and matrix decompositions^29^), rather than domain-specific microbiome feature engineering. To date, no modeling framework exists that systematically explores and optimizes microbial feature representation in concert with model class and hyperparameters, addressing all statistical properties of microbial nucleotide sequencing data^23,30^.

Here, we introduce *ritme* (= <u>R</u>obust <u>I</u>nference of Feature <u>T</u>ransformations and <u>M</u>odel <u>E</u>nsembles for Next-Generation Sequencing Data), a user-friendly and open-source software package that systematically optimizes microbial feature representations and ML pipelines tailored to the statistical properties of microbial sequencing data. *ritme* includes extensive and customizable feature-model-space configurations and state-of-the-art optimization algorithms to efficiently search the vast hyperparameter space on distributed compute environments. Its modular architecture makes it scalable to future additions of other multi-omics modalities. Beyond raw performance, *ritme* favors parsimonious feature-model configurations by selecting the simplest configuration that retains competitive predictive performance, yielding models that are easier to interpret and less prone to overfitting on the moderate sample sizes typical of microbiome studies. Accessibility across diverse programmatic skill sets is ensured with *ritme*’s multiple user interfaces and automated experiment tracking and evaluations.

We demonstrate the utility and versatility of *ritme* on three real-world case studies: (i) predicting an infant’s age from its gut microbiota, (ii) predicting ocean temperature from its microbiota, and (iii) classifying screen-relevant colorectal neoplasia (advanced adenoma or carcinoma vs. healthy colon) from human gut microbiota. In each use case, *ritme* improves predictive performance compared to original study methodologies while providing insights into the contribution and relevance of individual modeling hyper-parameters with regard to predictive performance. Together, these results underscore *ritme*’s potential to standardize the construction of robust predictive models from high-throughput sequencing data, which can subsequently serve as a starting point for further biological investigation.

## The *ritme* framework

### Overview of main functions in *ritme*

The main goal of *ritme* is to optimally infer the best feature representation and model selection for a specific predictive task from microbial nucleotide sequencing data. To do this it offers five main functionalities (Figure 1) that fully adhere to best practices in performing ML analyses of microbiome data^30^:

- `split_train -test` merges the provided feature and metadata tables and partitions the merged data into train and test sets. The split can be performed in a grouped manner — keeping related samples (e.g., from repeated measurements of the same host or field site) within a single partition. This enables controlled training designs, e.g., leave-one-subject out or leave-one-study out training routines. The split can additionally be stratified by one or more metadata columns to preserve their joint distribution across train and test, with stratification applied at the group level when grouping is active. split train test further supports two modeling regimes: a *static* regime in which a single time point per sample is used as input (**x***_t_ → y_t_*), and a *dynamic* regime in which the current snapshot is augmented with a fixed number (*N*) of preceding per-host snapshots to predict the target at time *t* (**x***_t−N_, …,* **x***_t_ → y_t_*).
- `find_best-model_config` performs a systematic, optimized exploration of the feature-model parameter space on the training set (see “Search for best feature-model configuration”), implementing CASH across both the microbiome-specific feature engineering (see “Microbiome-specific feature engineering”) and predictive model options (see “Predictive modeling frameworks”) for either regression or classification targets. It further provides insights into contributions of each feature representation and model component to predictive performance.
- `evaluate_tune_models` assesses the generalization capacity of optimal feature-model configurations per model type through performance evaluation on both training and independent held-out test sets, thereby mitigating selection bias and preventing inflated performance estimates characteristic of the “winner’s curse” phenomenon^23,31^.
- `explain_features` quantifies the contribution of individual features to the predictions of any selected best model on the held-out test set, using SHAP (SHapley Additive exPlanations) values^32^ for non-linear trainables and inferred coefficients for linear trainables.
- `explain_stability` assesses the robustness of these feature attributions across near-optimal models: it locates the best model’s top-ranked features in the importance rankings of comparably performing trials, matching features through their source features when trials differ in feature engineering.

**Figure 1.**
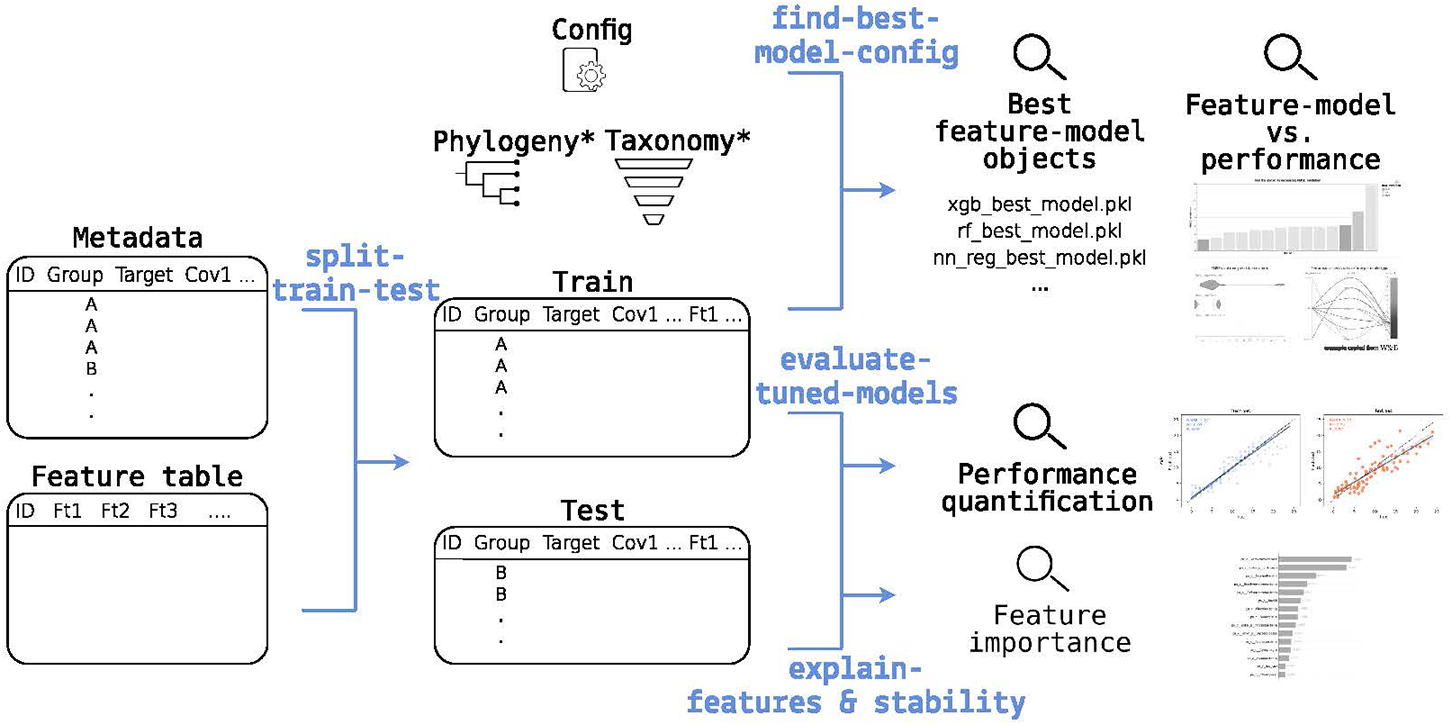
Overview of *ritme*’s data and modeling pipeline. `Split_train_test` merges the feature and metadata tables and partitions the data; `find_best_model_config` infers the optimal feature-model combination given an experiment configuration and provides insights into all explored combinations (inputs marked with an asterisk are optional); and `evaluate_tuned_models` quantifies predictive performance of the best feature-model combinations on the train and test sets. Feature contributions to the predictions of selected best models can be quantified with `explain_features`, and their robustness across comparably performing trials with `explain_stability`.

### Search for best feature-model configuration

*ritme*’s main function `find_best_model_config` optimally searches for the best feature-model combination on the provided dataset. Each combination is trained and evaluated by K-fold cross-validation on the provided training data (default *K* = 5), with folds partitioned according to the defined grouping and stratification criteria (see `split_train_test` in “Overview of main functions in *ritme*”). The *K* fold scores are aggregated per trial into a mean and standard error, providing both a point estimate of predictive performance and a measure of its uncertainty. Folds are trained in parallel for non-iterative learners and sequentially for iterative learners (Table S1). Combinations of feature engineering and model hyperparameters are iteratively evaluated per model type for a maximal run duration specified by the user (Figure 2).

**Figure 2.**
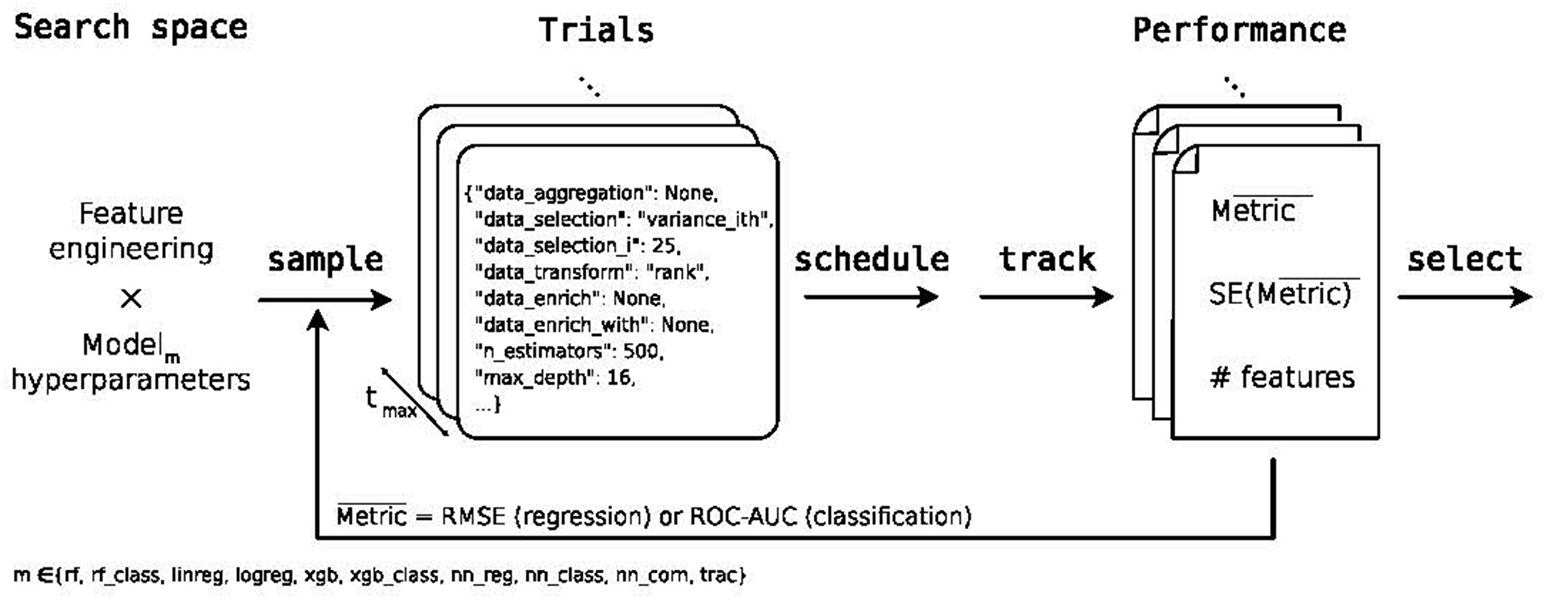
Inference of the optimal feature-model combination with find_best_model_config. Schematic of the search implemented in *ritme*’s `find_best_model_config` (built on Ray Tune). For each model class *m*, trials are launched for a specified maximal run duration *t*_max_. In each trial, Optuna samplers propose a joint configuration over the microbiome-specific feature-engineering and model-hyperparameter spaces. Scheduling and early stopping are handled by ASHA or HyperBand. Trial metrics are tracked with MLflow or Weights & Biases and computed by K-fold cross-validation on the training data (default *K* = 5; per-fold scores aggregated into a mean and standard error per trial). The best configuration within each model type is selected by the one-standard-error rule across all trials of that model type. RMSE = root mean squared error, ROC-AUC = area under the receiver operating characteristic curve, SE = standard error.

At the end of the search, the *one-standard-error rule* ^33^ is applied independently within each model type: among all trials of a given type whose mean cross-validation score lies within one standard error of that type’s best mean, the simplest is retained — yielding one winning configuration per model type and favoring parsimonious models over the nominal optimum. The simplicity criterion ranks configurations first by the count of retained features, and at equal feature counts by model-type-specific complexity measures (Table S2). The selected configuration is finally refit on the full training data and returned as a custom-made TunedModel object — encapsulating the chosen feature aggregation, selection, transformation, enrichment schema, and fitted estimator — thereby enabling reproducible, directly applicable prediction on any dataset with the same feature structure (Figure S1).

To perform the search for the best combined feature representation and modeling algorithm in a resource-aware manner, *ritme* leverages the Ray Tune framework^34^ enabling optimal computational resource management with parallelization across all available computational resources - from single workstations to high-performance computing environments. The Asynchronous Successive Halving Algorithm (ASHA)^35^ and HyperBand early stopping algorithm^36^ allow optimal scheduling of different hyperparameter configurations by terminating unpromising configurations early and adaptively allocating more resources to promising candidates, with the HyperBand algorithm providing deterministic scheduling when experiment reproducibility is needed (scheduling step in Figure 2).

To enable efficient hyperparameter search, hyperparameter suggestions are generated dynamically (define-by-run style) with different sampler algorithms using Optuna^37^ (sampling step in Figure 2), including the Tree-structured Parzen Estimator (TPE)^38,39^, the Covariance Matrix Adaptation Evolution Strategy (CMA-ES)^40,41^, a Gaussian process-based sampler (GPSampler)^37^, a Quasi Monte Carlo Sampler (QMCSampler) ^38,42^ and a random independent sampler^37^. For TPE, CMA-ES, and the GPSampler, the initial random warm-up scales with search-space complexity: max(20, 5 *d*_eff_) random configurations are drawn before model-guided sampling begins, where *d*_eff_ is the maximum number of hyperparameters the search space can propose in a single trial — providing enough seeds for the model-guided samplers on large spaces without wasting trials on small ones.

The predictive performance of different launched hyperparameter combinations is trackable with two popular ML experiment tracking tools (tracking step in Figure 2): MLflow^43^ and Weights and Biases^44^, for each of which a reproducible example template is provided for optimally evaluating *ritme* experiments (see Resource availability).

### Microbiome-specific feature engineering

*ritme* includes feature engineering methods that account for diverse statistical properties of microbial sequencing data (Figure 3). Feature representation is comprised of four sequential steps: taxonomic aggregation, feature selection, compositional data transformations, and feature enrichment. The possible combinations of these steps, along with their associated parameters (*i*, *q*, and *t*, Table S3) and model-specific parameters (see “Predictive modeling frameworks”) are systematically evaluated as hyperparameters during optimization.

**Figure 3.**
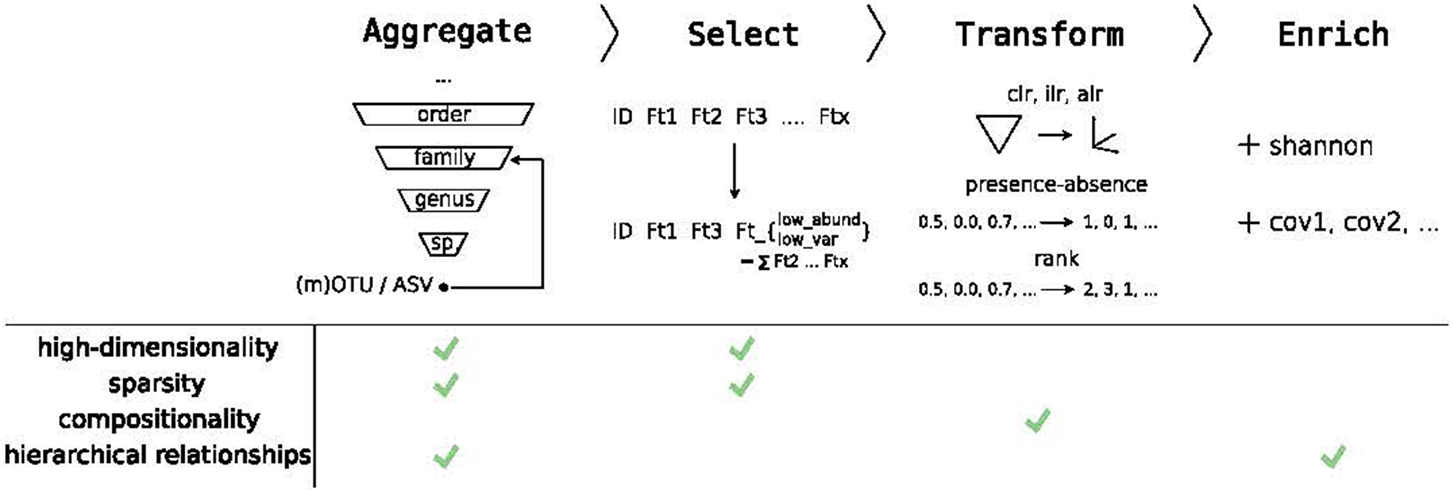
Four sequential microbiome-specific feature engineering steps implemented in *ritme*. Taxonomic aggregation, feature selection, compositional transformation, and feature enrichment are applied in sequence and together address the statistical properties of microbial sequencing data.

The taxonomic aggregation enables hierarchical aggregation of sequencing counts based on their taxonomic ranks where features are either left as-is (no aggregation) or summed according to rank. If the selected taxonomic entity is unknown for a specific sequence, the higher order rank is considered for aggregation.

To address the inherent sparsity and high-dimensionality of microbial sequencing data, *ritme* implements four parametric feature selection strategies based on either abundance or variance criteria (Table S3): top-rank selection (keep the top-*i* features), rank-referenced selection (keep features above the *i*-th-ranked reference), quantile selection (keep features above the *q*-th quantile), and absolute-threshold selection (keep features above a threshold value *t*). In every case the unselected features are summed into a single aggregate column to preserve the compositional nature of the data. Default hyperparameter ranges are inferred from the training data unless explicitly set by the user: *i* spans the integers 1*, …, p* (with *p* the number of features in the training set), *q* is explored over [0.5, 0.9], and *t* ranges from the minimum to the maximum observed feature-wise summed abundance or variance.

*ritme* addresses the compositionality constraint through several log-ratio transformations, including centered (clr), isometric (ilr) and additive (alr) log ratio^45–47^, a binary presence-absence transformation (pa)^48^ and rank order normalization (rank)^49,50^. For the additive log ratio transform, the feature with the highest relative abundance is selected as a reference.

Additionally, to allow for inclusion of contextual and microbiome-derived diversity information, *ritme* has the option to supplement the microbial features with Shannon diversity^51^ and/or metadata fields.

The resulting search space of feature engineering options and associated hyperparameters can be restricted by the user in case the biological interest lies in a subset of the available representations. Additionally, the user can define the starting point(s) of the hyperparameter search.

### Predictive modeling frameworks

*ritme* implements a diverse set of ML models spanning linear, ensemble, gradient-based and neural-network methods, and covers both parametric and non-parametric approaches (Tables 1, S4 and S5). Both regression and classification targets are supported, with each model class instantiated as a regressor or a classifier depending on the configured prediction task. Each model is evaluated on a validation set using the metric appropriate for its task — root mean squared error (RMSE) for regression and the area under the receiver operating characteristic curve (ROC-AUC) for classification (see “Search for best feature-model configuration”). For iterative gradient-based models, such as XGBoost and neural networks, training progress is monitored using MLflow^43^ or Weights and Biases^44^, with scheduler’s early stopping mechanisms in place to optimize search efficiency (see “Search for best feature-model configuration”).

Beyond the standard regression and classification tasks, *ritme* additionally supports an ordinal regression task via the CORN neural network^52^ (nn corn), which is particularly suited for targets with a natural class order. Trained ordinal regression (nn corn) neural networks can be evaluated under either regression or classification metrics (Table 1).

**Table 1.** Predictive models implemented in *ritme*. The Task column indicates the supported prediction tasks: R = regression, C = classification. MSE = mean squared error, trac = tree-aggregation of compositional data, CORN = Conditional Ordinal Regression for Neural networks.

| Name | Algorithm | Implementation | Training Loss | Task |
| --- | --- | --- | --- | --- |
| linreg | Elastic net linear regression | scikit-learn<sup>77</sup> | Elastic Net | R |
| rf | Random forest | scikit-learn | MSE | R |
| trac | trac<sup>53</sup> | adapted from c-lasso<sup>78</sup> | Squared error | R |
| xgb | XGBoost<sup>79</sup> | XGBoost<sup>79</sup> | Squared error | R |
| nn_reg | Neural network | PyTorch Lightning<sup>80</sup> | MSE | R |
| nn_corn | CORN<sup>52</sup> | coral-pytorch<sup>52</sup> | CORN loss<sup>52</sup> | R, C |
| nn_class | Neural network | PyTorch Lightning | Cross-entropy | C |
| logreg | Logistic regression | scikit-learn | Cross-entropy | C |
| rf_class | Random forest | scikit-learn | Gini impurity | C |
| xgb_class | XGBoost<sup>79</sup> | XGBoost<sup>79</sup> | Cross-entropy | C |

Each model is integrated with the feature engineering pipeline described in “Microbiome-specific feature engineering”, facilitating systematic evaluation across diverse feature representation for each model. Only the trac model^53^ contains a complementary approach by directly incorporating phylogenetic hierarchies and dynamically learning optimal taxonomic aggregation levels during training. Prior to modeling, features for linear and logistic regression are standardized using standard scaling, while neural networks utilize batch normalization^54^ to normalize features before they are passed through the network. Similar to feature engineering hyperparameters, users can restrict the model hyperparameters tested and define the starting point(s) of the search.

### Implementation and availability

*ritme* is implemented as a modular open-source Python 3 package, offering a user-friendly interface through both a Python API and a command-line interface, facilitated by the *Typer* library^55^. Its modular architecture facilitates future extensions with additional feature representation or modeling algorithms, enabling the open-source community to incorporate feature representations of other multi-omics modalities and domain-specific models. Furthermore, each search is fully specified by a single experiment configuration file — encompassing the searched feature and model spaces, optimization settings, and random seeds — which is persisted alongside every experiment, thereby enabling the reproduction of previously conducted analyses through the exchange of this file alone. To ensure code reliability and robustness, *ritme* was developed in accordance with best practices in software development^56,57^, incorporating unit testing and continuous integration on Ubuntu and macOS. *ritme* is openly available together with comprehensive tutorials and reproducible usage examples (see Resource availability).

## Results

### *ritme* outperforms the original studies with more parsimonious feature-model configurations

We evaluated *ritme* on three real-world datasets previously used to predict a target from microbial features (Table 2): infant age (use case 1) and ocean temperature (use case 2) as regression tasks, and screen-relevant colorectal neoplasia (use case 3) as a binary classification task. We split each dataset into a training set (80% of samples) and a held-out test set (20%), ran *ritme*’s search on the training set for 23 hours per model type, and reproduced the original studies’ feature engineering and modeling pipelines on the same split. For *ritme*, we report the one-standard-error winner with the highest mean cross-validation score across model types. For all three use cases, the optimal feature-model configuration identified by *ritme* outperforms the original study configurations when evaluated on the same held-out test set (Tables 3 to 5 and figs. S2 to S4). In use cases 2 and 3, *ritme*’s best model also showed a substantially smaller training-test gap than the original model setup (Figures S3 and S4).

**Table 2.** Datasets used for use cases comparing *ritme* to original models. Number of features (# features) include features that are non-zero in at least one sample. U1 and U2 are regression tasks; U3 is a binary classification task. R = regression, C = classification, HTS = high-throughput sequencing, Ref. = reference, OTUs = operational taxonomic units, miTAGs = metagenomic 16S ribosomal RNA gene tags, SRN = screen-relevant neo-plasia (advanced adenoma or carcinoma).

| Use case | Target | # Samples | # Features | Feature type | HTS method | Ref. |
| --- | --- | --- | --- | --- | --- | --- |
| U1 | Infant age (months, R) | 448 | 18,150 | 16S OTUs<sup>17,18</sup> | Amplicon | Subramanian et al.<sup>66</sup> |
| U2 | Ocean temperature (celsius, R) | 139 | 35,651 | 16S OTUs derived from miTAGs<sup>19,73</sup> | Shotgun | Sunagawa et al.<sup>8</sup> |
| U3 | SRN (binary, C) | 489 | 11,282 | 16S OTUs | Amplicon | Topçuoğlu et al.<sup>75</sup> |

**Table 3.** Use case 1: held-out test performance and configuration components of the original published approach and *ritme*. Performance metrics and configuration components of the original approach^66^ and *ritme*, reported both for a no-metadata ablation (microbial features and Shannon diversity only; external metadata excluded from the search) and the full *ritme* search. For the original feature selection, the threshold is based on relative abundance values. Metadata used for feature enrichment with *ritme* includes the following variables at sampling: health status, milk diet, weaning status and antibiotics exposure 7 days prior. Best performance per metric in boldface. seq = sequences, RMSE = root mean squared error, R^2^ = coefficient of determination.

|  |  | **Original** | *ritme* (no metadata) | *ritme* |
| --- | --- | --- | --- | --- |
| **Metrics** | RMSE | 3.498 | 3.183 | **2.892** |
|  | R<sup>2</sup> | 0.669 | 0.726 | **0.773** |
| **Config** | Algorithm | Random Forest<sup>77</sup> | **xgb** | **xgb** |
|  | Aggregation | - | - | - |
| | Selection | $\geq 10^{-3}$ in $\geq 2$ samples | variance_threshold,<br>$t = 10^{-6}$ | variance_threshold,<br>$t = 10^{-6}$ |
|  | Transform | Rarefaction to 2,000 seq/sample | ilr | ilr |
|  | Enrichment<br># Features | -<br>850 | shannon<br>205 | metadata_only<br>215 |

Consistently, *ritme*’s best feature-model configurations differed from the original approaches. As for the model algorithm, *ritme*’s selected model type overlapped with the original set-up only in use case 2 (both ElasticNet linear regression). In terms of feature engineering, *ritme* identified more parsimonious configurations with fewer features than the original set-ups in all use cases. Both modeling set-ups used the most granular microbial feature count representation, with no taxonomic aggregation selected in any use case. Feature selection criteria aligned only in use case 2, where both set-ups retained features by abundance; in use case 1, *ritme* substituted a variance-based criterion for the original relative-abundance threshold, and in use case 3 it introduced abundance-based selection where the original applied none. Transformation choices diverged substantially only in use case 1, where *ritme* selected the compositionality-aware ilr transform over the original rarefaction; in use cases 2 and 3, both set-ups relied on comparable generic rescalings (Tables 4 and 5).

**Table 4.** Use case 2: held-out test performance and configuration components of the original published approach and *ritme*. Performance metrics and configuration components of the original approach^8^ and *ritme*, reported both for a no-metadata ablation (microbial features and Shannon diversity only; external metadata excluded from the search) and the full *ritme* search. For the original feature selection, the threshold is based on relative abundance values. *x*0 represents a pseudocount = 1e-6. Metadata used for feature enrichment with *ritme* includes sampling depth, environmental feature, latitude, and longitude. Standard scaling denotes the linear-model-specific transformation step that *ritme* applies to all features of linreg and logreg models prior to fitting, in addition to the searched transform (see “Predictive modeling frameworks”). Best performance per metric in boldface. CV = cross-validation, RMSE = root mean squared error, R^2^ = coefficient of determination.

|  |  | Original | *ritme* (no metadata) | *ritme* |
| --- | --- | --- | --- | --- |
| **Metrics** | RMSE | 2.011 | 1.580 | **1.435** |
| | $R^2$ | 0.915 | 0.948 | **0.957** |
| **Config** | Algorithm | Elastic Net<sup>77</sup> with 5-fold CV | `linreg` | `linreg` |
|  | Aggregation | - | - | - |
| | Selection | $\geq 10^{-4}$ in any sample | - | `abundance_topi`, $i = 1,971$ |
| | Transform | $\log_{10}(x + x_0)$ & rank | standard scaling & rank | standard scaling & rank |
|  | Enrichment | - | - | `shannon_and_metadata` |
|  | # Features | 7,250 | 1,668 | 1,981 |

**Table 5.**
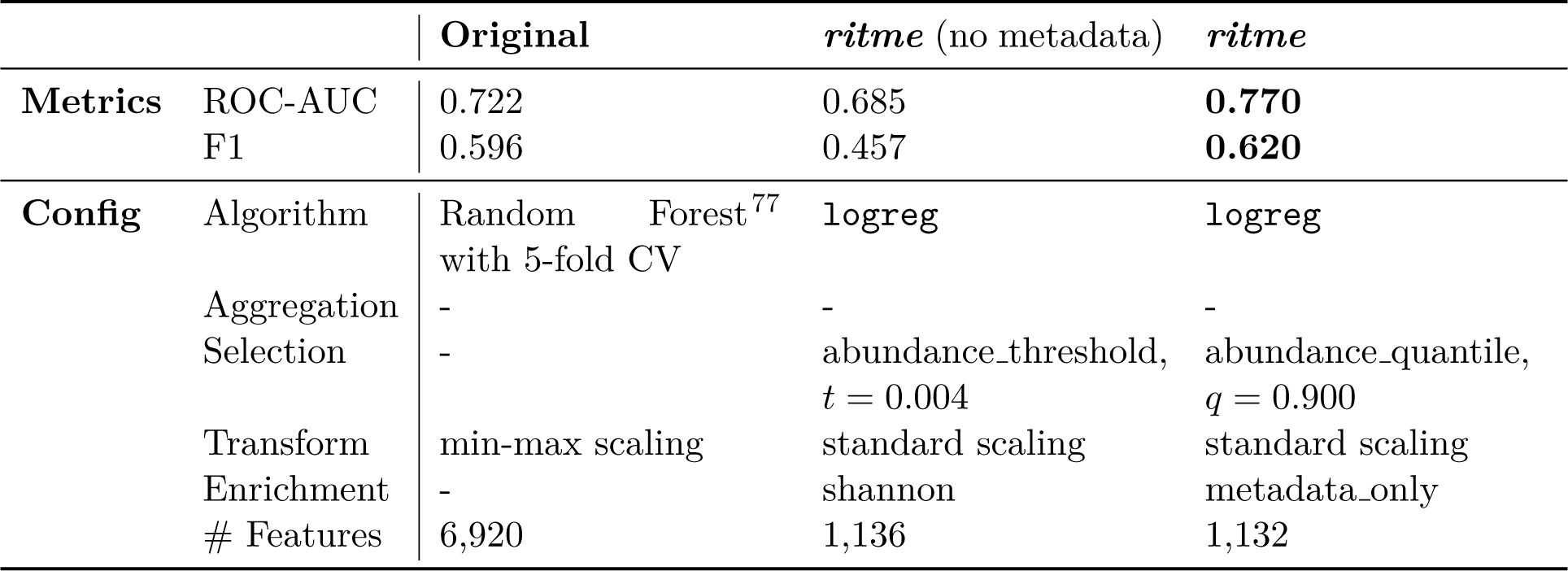
Use case 3: held-out test performance and configuration components of the original published approach and *ritme*. Performance metrics and configuration components of the original approach^75^ and *ritme*, reported both for a no-metadata ablation (microbial features and Shannon diversity only; external metadata excluded from the search) and the full *ritme* search. Metadata used for feature enrichment with *ritme* includes age, gender, and BMI. Standard scaling denotes the linear-model-specific transformation step that *ritme* applies to all features of linreg and logreg models prior to fitting (see “Predictive modeling frameworks”); no transformation was selected within the searched transform space. Best performance per metric in boldface. ROC-AUC = area under the receiver operating characteristic curve, F1 = macro-averaged F1 score, CV = cross-validation.

|  |  | **Original** | *ritme* (no metadata) | *ritme* |
| --- | --- | --- | --- | --- |
| **Metrics** | ROC-AUC | 0.722 | 0.685 | **0.770** |
|  | F1 | 0.596 | 0.457 | **0.620** |
| **Config** | Algorithm | Random Forest<sup>77</sup><br>with 5-fold CV | **logreg** | **logreg** |
|  | Aggregation | - | - | - |
| | Selection | - | abundance_threshold,<br>$t = 0.004$ | abundance_quantile,<br>$q = 0.900$ |
|  | Transform | min-max scaling | standard scaling | standard scaling |
|  | Enrichment | - | shannon | metadata_only |
|  | # Features | 6,920 | 1,136 | 1,132 |

In all three use cases, *ritme* identified that enriching microbial features with metadata — alone (`metadata_only`, use cases 1 and 3) or together with Shannon diversity (`shannon_and_metadata`, use case 2) — improved predictive performance beyond what was employed in the original studies. Notably, even when non-microbial metadata covariates are excluded from the search (no-metadata ablation for the winning model type under the same split; see Methods), *ritme* still outperforms the original approaches in two of three use cases (U1 RMSE 3.18 vs. 3.50; U2 1.58 vs. 2.01; no-metadata columns in Tables 3 and 4), confirming that *ritme*’s feature-model search alone identifies stronger microbiome-only configurations. In use case 3, by contrast, the no-metadata ablation reached a ROC-AUC of 0.69 and a macro-F1 of 0.46 (vs. 0.72 and 0.60 for the original; no-metadata column in Table 5). A dedicated comparison of the two *ritme* models (Figure S5) shows that the metadata covariates leave the microbial signature essentially unchanged (13 of the 15 top-ranked features shared, Spearman’s *ρ* = 0.95 between absolute coefficients); their benefit is instead confined to one of the two colonic lesion types pooled in the prediction target: microbial features discriminated carcinomas from normal colons, whereas the precancerous lesions were discriminated by age alone. The residual deficit of the no-metadata *ritme* model relative to the original set-up reflects the model family rather than the feature engineering, as a random forest fitted on *ritme*’s engineered features attained a ROC-AUC of 0.746 on the same split.

### *ritme* provides insights into optimal feature representation and model selection

During inference, each feature-model configuration proposed by *ritme*’s search is evaluated by K-fold cross-validation on the training data, with the K-fold scores summarized by their per-trial mean (“Search for best feature-model configuration” and Figure 2). With *ritme*, users can inspect these configurations and their contributions to the predictive performance (Figure 1). Examining configurations across the three use cases (Figure 4) shows that cross-validation performance distributions vary by feature engineering approach and model type, although the displayed configurations also vary in model-specific hyperparameters within each category, which can confound the observed between-category differences. Notably, the top-performing configuration identified for each use case (Tables 3 to 5) does not consistently represent the best median performance across trials, reflecting the multiplicity of near-optimal models — the *Rashomon effect* — whereby many distinct feature-model configurations attain comparable predictive accuracy^58,59^.

**Figure 4.**
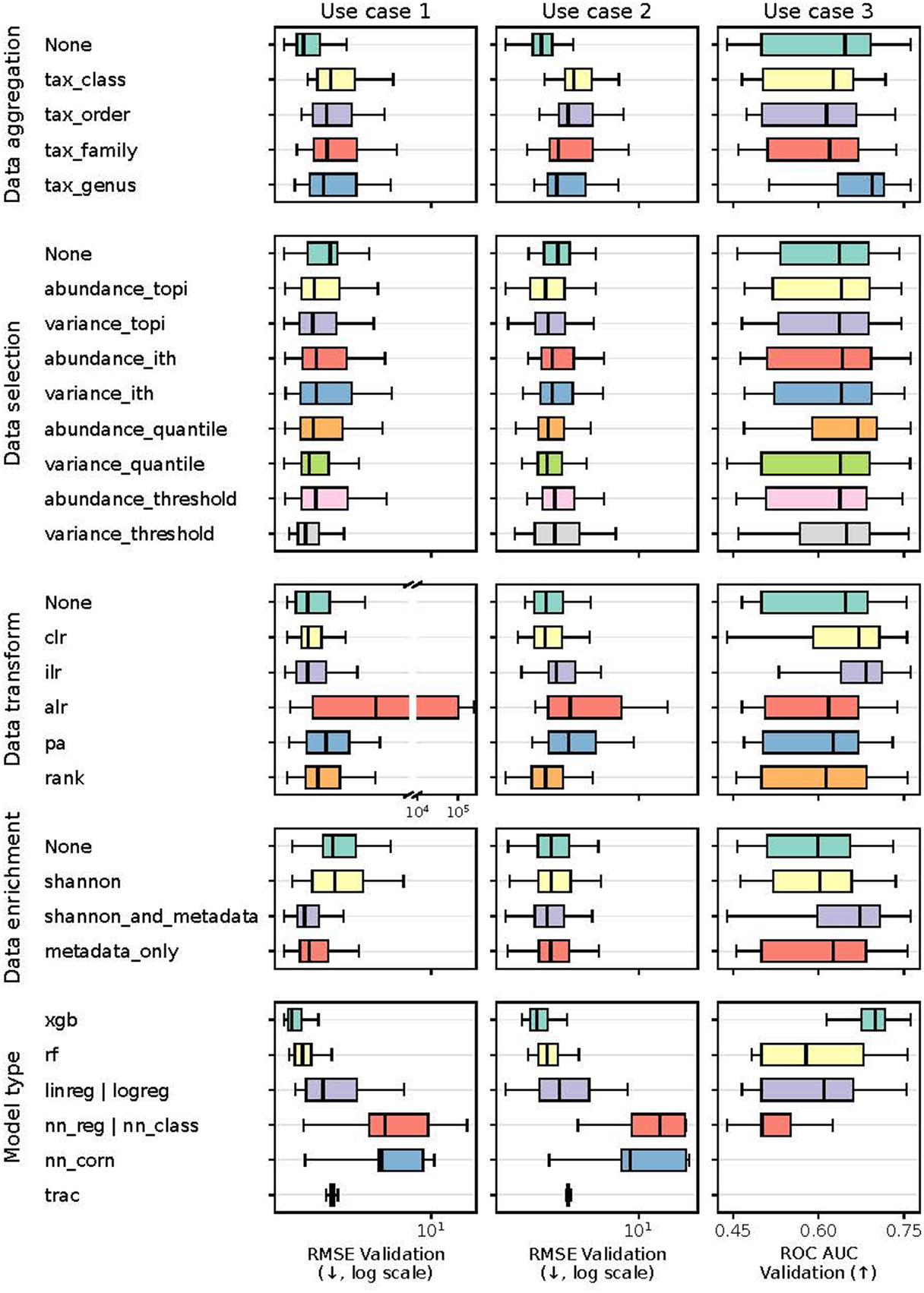
Cross-validation performance across tested feature engineering and model configurations. Distributions of per-trial cross-validation performance across *ritme*’s tested feature engineering and model configurations for all three use cases: validation root mean squared error (RMSE) for the regression use cases 1 and 2 and validation area under the receiver operating characteristic curve (ROC AUC) for the classification use case 3. Each trial contributes the mean of its K validation-fold scores.

This multiplicity of near-optimal models also bears on interpretation, as such configurations need not agree on which features matter. *ritme*’s `explain_stability` therefore quantifies, for the task at hand, how well the top-ranked features of the selected best model recur across comparably performing trials (see “Overview of main functions in *ritme*”). The three use cases span the spectrum of task-specific outcomes this diagnostic reveals (Figures S6 to S8): stable feature importances that support downstream biological investigation (use case 3 Figure S8), attributions spread across many interchangeable feature sets, directing interpretation to the ensemble of near-optimal models rather than any single one (use case 2 Figure S7), and a representation effect in which trial-specific ilr coordinates preclude cross-trial comparison of microbial features (use case 1 Figure S6).

Relating the number of features retained by the tested configurations (Figure 5a) to their predictive performance (Figure 5b) further allows users to identify the model family or individual trial that achieves acceptable performance with the fewest features, supporting parsimonious model choices for downstream investigation.

**Figure 5.**
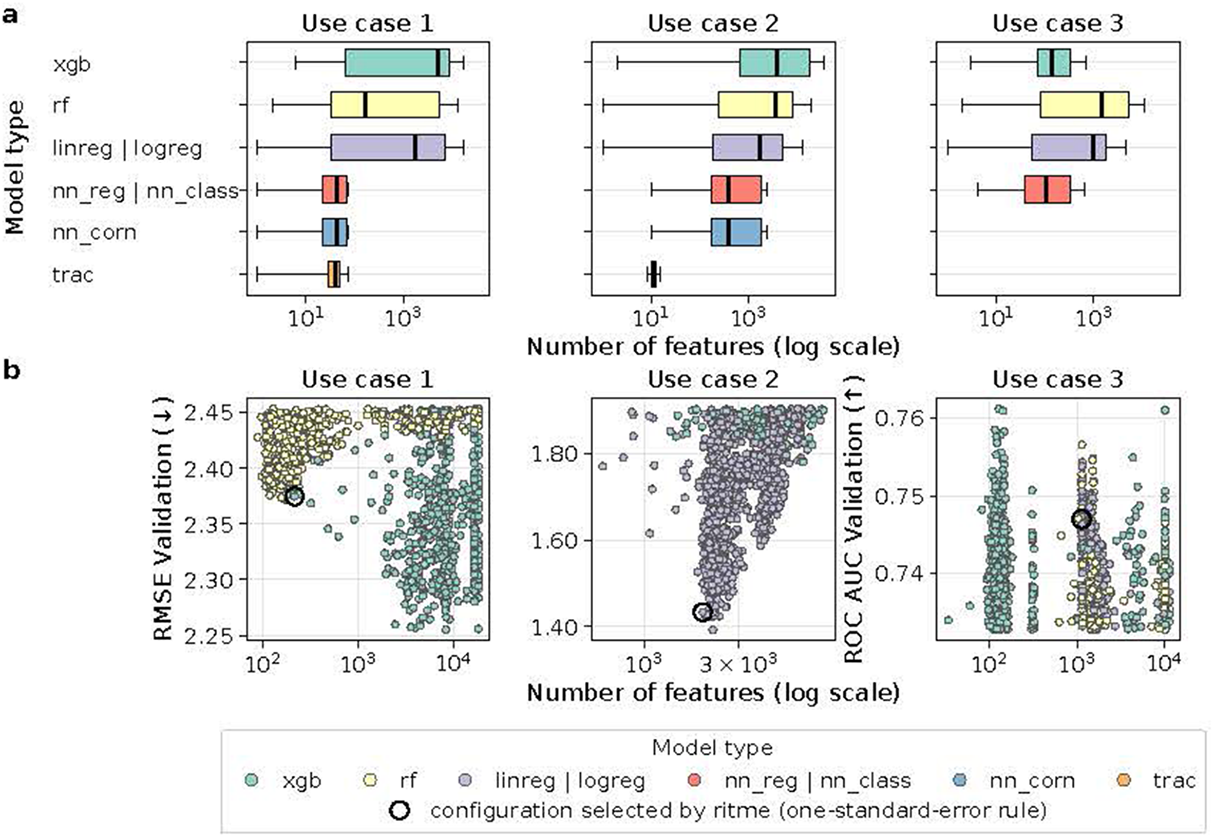
Number of retained features across tested configurations and its relation to validation performance. Number of features retained by the tested configurations for all three use cases, (**a**) grouped by model type and (**b**) related to validation performance (per-trial mean across K validation folds; RMSE for regression and ROC-AUC for classification) for the 1,000 best-performing trials of each use case. Each point in (b) represents one trial, colored by model type; the circled point marks the configuration selected by *ritme*’s one-standard-error rule. Regression and classification counterparts of a model class share a row (e.g. `linreg | logreg`).

Analysis of cross-validation performance trajectories across trials reveals model-specific search effectiveness (Figure 6). Models with sufficient completed trials — notably `xgb` and `trac` — showed clear performance gains as TPE-guided sampling took over from the initial adaptive random warm-up phase (see “Search for best feature-model configuration”). A dedicated sampler benchmark on use case 1 confirms this: TPE reached a lower validation RMSE than purely random search within the first hour and retained this advantage over a 12-hour budget (Figure S9). Linear models (linreg, logreg) showed limited additional gains beyond the warm-up, reflecting their compact hyperparameter space. Neural networks, with their longer per-trial training cost, completed substantially fewer trials (*≤* 190 per use case) and therefore remained close to their warm-up baseline. For these compute-heavy models, constraining hyperparameter ranges to smaller architectures may improve search efficiency. Users can experiment with alternative search algorithms through *ritme* to optimize exploration-exploitation balance for their specific datasets.

**Figure 6.**
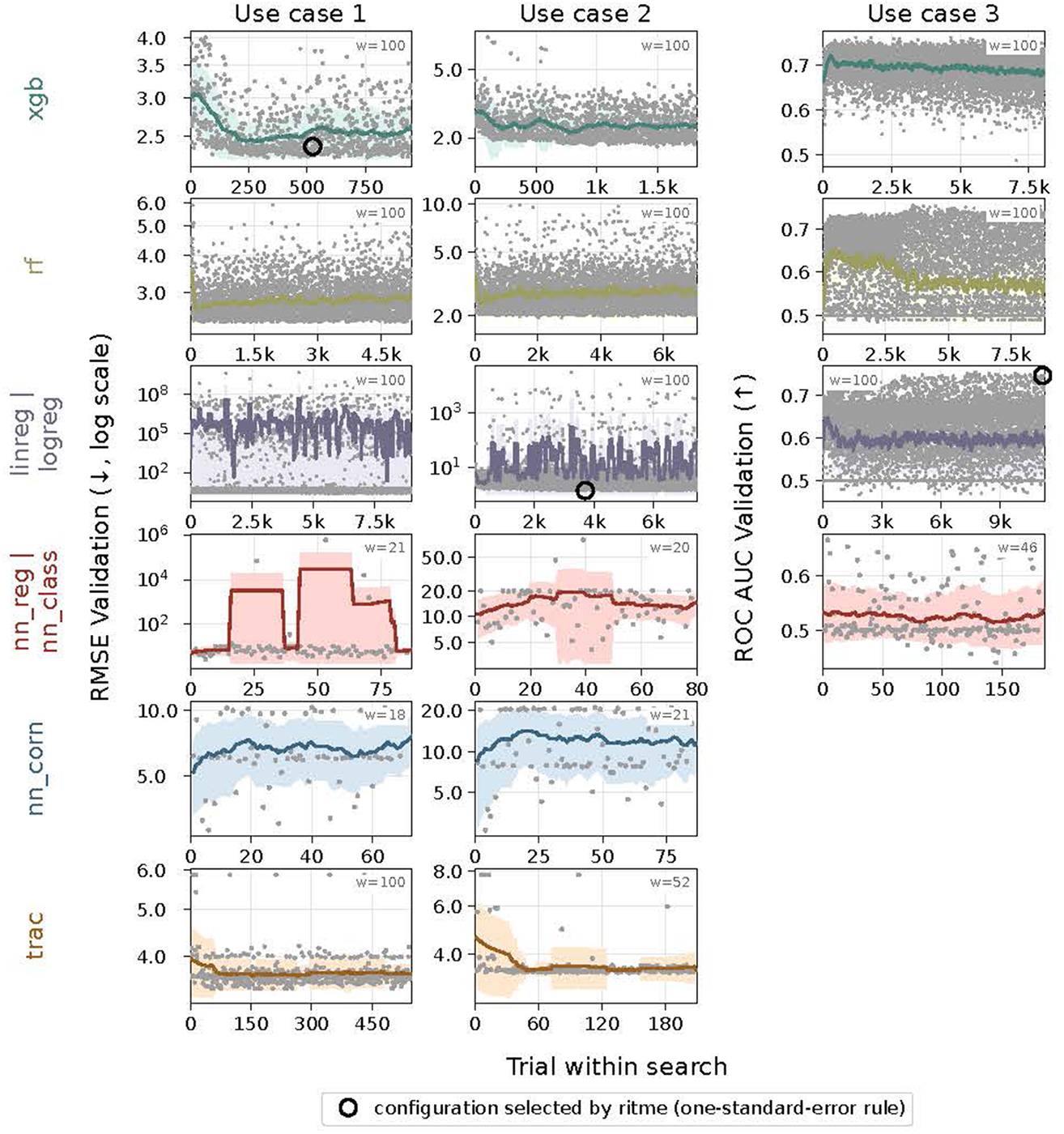
Validation-performance trajectories across trials per model type. Validation-performance trajectories across trials for all three use cases (per-trial mean across K validation folds; RMSE for regression and ROC-AUC for classification), shown per model type. Each gray dot represents one trial, colored lines the rolling mean and shaded areas its *±*1 standard deviation, with the window size adapting to the number of completed trials (reported as w in each panel). The circled point marks the configuration selected by *ritme*’s one-standard-error rule; panels are empty where a model class does not apply to the task.

### Microbiome-specific engineering enables *ritme* to outperform generic AutoML

To test whether microbiome-specific feature engineering improves predictive performance over generic preprocessing, we compared *ritme* with three AutoML baselines that likewise optimize preprocessing jointly with model class and hyperparameters, but over generic operations only (Table S6). The frameworks *auto-sklearn* and *TPOT* were run under matched compute budgets; the classification pipeline *mAML* was run as published (see Methods). On the primary task metric, *ritme* achieved better held-out test performance in six of the seven baseline comparisons (Table 6): it surpassed *auto-sklearn* in all three use cases, *mAML* in the classification use case, and *TPOT* in use cases 2 and 3, with *TPOT* in use case 1 remaining the sole exception (Figures S2 to S4). A compute-scaling benchmark on use case 1 (4 to 128 CPU cores) further demonstrated *ritme*’s compute efficiency: from 32 allocated cores onwards, *ritme* reached the lowest validation RMSE of all frameworks while sustaining high CPU utilization across every allocation, whereas *TPOT* left a growing share of the larger allocations idle (Figure S10). These results indicate that microbiome-specific feature engineering contributes to inferring predictive models from microbial feature count tables, while also showing that a well-tuned generic pipeline optimizer can match *ritme* on individual datasets.

**Table 6.** Predictive performance of *ritme* and the AutoML baselines on held-out test sets across the three use cases. Use cases 1 and 2 are regression tasks (reported as RMSE and R^2^); use case 3 is a classification task (reported as ROC-AUC and macro-F1). Top performance per metric in boldface, second-best underlined. *mAML* implements only classification models and was therefore not run on the regression use cases 1 and 2, indicated by “-”. RMSE = root mean squared error, R^2^ = coefficient of determination, ROC-AUC = area under the receiver operating characteristic curve, F1 = macro-averaged F1 score.

| Use case | Metric | *ritme* | *auto-sklearn* | *TPOT* | *mAML* |
| --- | --- | --- | --- | --- | --- |
| **1** | RMSE | <u>2.892</u> | 2.960 | **2.826** | - |
| | $R^2$ | <u>0.773</u> | 0.763 | **0.784** | - |
| **2** | RMSE | **1.435** | 2.048 | <u>1.906</u> | - |
| | $R^2$ | **0.957** | 0.912 | <u>0.924</u> | - |
| **3** | ROC-AUC | **0.770** | 0.742 | <u>0.745</u> | 0.632 |
|  | F1 | <u>0.620</u> | 0.434 | **0.708** | 0.616 |

## Discussion

*ritme* provides a domain-adapted framework for optimized predictive modeling on microbial sequencing data that addresses key statistical properties of these datasets — sparsity, high dimensionality, compositionality, and hierarchical structure — while jointly optimizing feature representation and model class.

Across three independent use cases, *ritme* outperformed original study modeling pipelines (Tables 3 to 5) and, on the primary task metric, the AutoML baselines in six of seven comparisons (Table 6). These gains stem from *ritme*’s central methodological novelty, the joint optimization of microbiome-specific feature representations with model class and hyperparameters, rather than from a new predictive algorithm: the model space comprises methods already in common use in the field (Table 1). The framework is therefore agnostic to the estimator employed, and its modular design allows further algorithms to be added as the field adopts them.

The comparison against the AutoML baselines emphasizes that domain knowledge matters for predictive performance, with *ritme*’s microbiome-specific feature engineering options generally outperforming the exclusively generic preprocessing spaces searched by these baselines (Table S6). That *TPOT* achieved a lower test RMSE in use case 1 indicates that on an individual dataset a sufficiently thorough generic search can match a domain-specific one in predictive accuracy. Predictive accuracy is, however, only one axis of comparison: unlike the AutoML baselines, *ritme* couples its search with transparent diagnostics of how feature and model choices shape performance (Figures 4 to 6) and how stable the top-ranked features are across near-optimal models (Figures S6 to S8).

Parsimony is another such axis: driven by the one-standard-error selection rule (see “Search for best feature-model configuration”), *ritme* identified feature-model configurations that retained substantially fewer features than the original set-ups in all use cases. Parsimonious configurations support interpretability and lower the risk of overfitting on the moderate sample sizes typical of microbiome studies. At the same time, automated optimization operates strictly within its predefined configuration space: known problem structure beyond this space — such as the repeated sampling of individual infants in use case 1, which *ritme* accounts for only through grouped data splitting (see Methods) — is not exploited by the search, and a bespoke model built around such structure may achieve higher predictive accuracy at the cost of greater development effort. In practice, users can also inject their own expertise by constraining the search to biologically plausible subspaces or by seeding initial configurations (“Microbiome-specific feature engineering” and “Predictive modeling frameworks”). Moreover, while *ritme* is optimized for microbiome sequencing data, it can be applied to any 2D multi-omics feature count table by deselecting non-applicable feature engineering steps (e.g., taxonomic aggregation or compositional transforms).

The recurrence of parsimonious configurations that retain competitive predictive performance is consistent with the *Rashomon-set* perspective on machine learning: for a given problem there often exists a large set of near-equally-accurate models, and simple-yet-accurate models tend to be found within it^58,60^. *ritme*’s one-standard-error rule operationalizes this perspective: among the near-optimal trials — those whose mean cross-validation score lies within one standard error of the best — it retains the simplest configuration (see “Search for best feature-model configuration”), thereby committing to a parsimonious member of this set rather than to the single highest-scoring trial. Relatedly, because near-optimal models can rely on different feature subsets, the variables highlighted by any single best model represent one of several plausible explanations rather than a unique one^59^. *ritme* makes this multiplicity inspectable by logging the full set of near-optimal configurations alongside their feature representations and complexity (Figures 4 and 5), and its `explain_stability` function quantifies the consequences for interpretation directly: it reveals, per predictive task, whether the best model’s top-ranked features recur across comparably performing trials and hence whether they warrant downstream biological interpretation (Figures S6 to S8).

As an open-source software built in a modular fashion, we plan different future extensions to *ritme*. Automatically selecting and adapting the hyper-parameter search algorithm to the given data’s search space characteristics will enable even more data-driven inference of the optimal feature-model configuration. Also, instead of optimizing feature-model combinations per model algorithm, a joint optimization will be evaluated. Functional-pathway aggregation will be added as a parallel feature-representation axis to complement the current taxonomic-aggregation hierarchy. Finally, we plan to couple *ritme*’s programmatic search with autonomous large language model agents that iteratively propose, run, and evaluate feature-model experiments, advancing toward agent-guided experimentation^61^.

In summary, *ritme* offers a data-driven, compute-efficient framework that standardizes the joint optimization of feature representations and models for microbiome sequencing data. By combining domain-specific feature engineering with optimized hyperparameter search and transparent experiment tracking, it delivers superior predictive performance with parsimonious feature-model configurations and actionable insight into which biological and statistical choices drive that performance. By replacing subjective, *ad hoc* modeling decisions with systematic optimization, *ritme* enables researchers to identify best-performing yet parsimonious feature-model combinations from microbiome sequencing data in a standardized manner. We anticipate that *ritme* will help mitigate reproducibility concerns in microbiome research^62^ by providing robust predictive models that can serve as a starting point for downstream biological investigation.

## Methods

### Comparative modeling

#### *ritme* vs. original studies

To demonstrate the capabilities of *ritme*, we selected three real-world datasets which were all previously used to predict a target (continuous in use cases 1 and 2; binary in use case 3) from microbial features (Table 2). For each use case, we initially split the data using the *ritme* `split_train_test` method into a train set and a test set, 80% and 20% of all samples. For use case 1, which includes repeated measurements per host, we applied a grouped split by host identifier to prevent information leakage between train and test sets. On the train set, we launched *ritme*’s `find_best_model_config` and additionally reproduced the original publication’s feature engineering and modeling set-ups for comparison. The left-out test set was used to evaluate the predictive performance of the original approaches and the best feature-model configurations selected by *ritme* via the one-standard-error rule on the K-fold cross-validation scores computed on the training set (see “Search for best feature-model configuration”). For downstream reporting per use case, we then selected — across the per-model-type 1-SE winners — the trial with the highest mean cross-validation score.

For all use cases, *ritme*’s find best model config was launched for a duration of 23 hours per model type (Table 1). For use cases 1 and 2 (regression), all regression-compatible model types were included. For use case 3 (binary classification), the classification model types were considered. The hyperparameter search spaces for all use cases were searched using the TPE-Sampler.

To isolate *ritme*’s microbial feature-model contribution from external-metadata signal, we additionally ran a no-metadata ablation per use case (external metadata excluded from the search; Shannon diversity still allowed) for the winning model type, under the same train/test split and one-standard-error selection protocol.

#### *ritme* vs. AutoML baselines

To test whether microbiome-specific data engineering actually improves predictive performance, we compared *ritme* against three AutoML baselines on the same training data and the same held-out test sets. The baselines were selected among frameworks applicable to supervised prediction from microbial feature count tables (Table S6) that jointly optimize feature preprocessing together with model class and hyperparameters (CASH): the general-purpose AutoML systems *auto-sklearn* ^29^ and *TPOT* ^26^, and *mAML* ^24^, an AutoML pipeline designed specifically for microbiome-based classification tasks. Notably, although all three baselines search feature preprocessing jointly with model hyperparameters, their searched preprocessing spaces comprise exclusively generic operations; *ritme* remains, to our knowledge, the first framework to include microbiome-specific feature engineering options within the CASH search itself (Table S6).

*auto-sklearn* and *TPOT* were compared under a matched set-up: both were launched for the same run duration and with the same computational resources as *ritme*, restricted to the estimator matching the best-performing model type identified by *ritme* (e.g., for xgb – *auto-sklearn*’s `gradient boosting` and *TPOT*’s `XGBRegressor`; for `linreg` – *auto-sklearn*’s sgd and *TPOT*’s `ElasticNetCV)`, and both optimized the primary task metric (RMSE for the regression use cases U1 and U2; ROC-AUC for the classification use case U3). *auto-sklearn* was additionally constrained to return a single best model rather than an ensemble. *mAML*, by contrast, implements a fixed grid of feature scalers and classifiers rather than a budgeted search and was therefore run as published: it neither consumes the shared time budget nor selects on the primary task metric, and it applies to the classification use case (U3) only.

### Data fetching and processing

#### Use case 1 - infant age

Sequencing data and associated metadata were fetched with *q2-fondue* ^63^ from the NCBI Sequence Read Archive (SRA)^64^ using the reported BioProject ID PRJEB5482, and analyzed downstream using QIIME 2^65^. Additional metadata were retrieved from the publication’s^66^ supplementary materials. To replicate the analysis described in the original publication, forward and reverse reads were trimmed to at most 162 nucleotides in length using *q2-cutadapt* ^67^. The overlapped paired reads were clustered sharing *≥* 97% similarity matched to the 13 8 99% Greengenes reference^68^ using *q2-vsearch* ^69^, with remaining reads clustered *de novo*. The resulting feature count table consists of 448 samples from 25 unique infants with 18,150 16S OTUs (operational taxonomic units) (Table 2).

Taxonomy was assigned by a trained naive Bayes classifier on the Green-genes 13 8 99% reference database using *q2-feature-classifier* ^70^, after extracting the V4 region. The phylogenetic tree was inferred by aligning sequences with MAFFT and masking the alignment using *q2-alignment* ^71^, constructing and midpoint-rooting the tree using FastTree in *q2-phylogeny* ^72^.

#### Use case 2 - ocean temperature

Sequencing data, associated metadata, and sequence taxonomy were fetched from the original publication’s^8^ companion website https://ocean-microbiome.embl.de/companion.html. The resulting sequence feature count table consists of 139 samples with 35,651 16S OTUs, derived from metagenomic 16S ribosomal RNA gene tags (miTAGs)^19,73^ and clustered at 97% similarity to the release 115 SILVA 16S reference database^74^ (Table 2).

The phylogenetic tree was inferred by aligning sequences with MAFFT and masking the alignment using *q2-alignment* ^71^, constructing and midpoint-rooting the tree using FastTree in *q2-phylogeny* ^72^.

#### Use case 3 - colorectal neoplasia classification

Sequencing data, associated metadata, and sequence taxonomy were fetched from the original publication’s^75^ companion GitHub repository at https://github.com/SchlossLab/Sze_CRCMetaAnalysis_mBio_2018. The resulting sequence feature count table consists of 489 samples with 11,282 16S OTUs of the V4 region clustered at 97% similarity (Table 2). The binary target srn (screen-relevant neoplasia: advanced adenoma or carcinoma vs. non-advanced adenoma, normal, or high-risk-normal colon) was derived from the metadata per the original publication’s definition.

#### Use of AI-assisted tools

GitHub Copilot and Claude Code were used to assist with software testing, debugging, documentation, and data analysis. All AI-assisted outputs were reviewed by the authors.

## Resource availability

### Lead contact

Requests for further information and resources should be directed to and will be fulfilled by the lead contact, Nicholas A. Bokulich (nicholas.bokulich@ hest.ethz.ch).

### Materials availability

*ritme* is installable via Anaconda at https://anaconda.org/adamova/ritme.

### Data and code availability

**Software:** *ritme* is open-source under the BSD 3-clause license, available at https://github.com/adamovanja/ritme and archived on Zenodo^76^ at https://zenodo.org/records/17537117. Reproducible examples for the Python API and command-line interface, together with templates for evaluating *ritme* experiments with MLflow or Weights & Biases, are provided in the repository. **Data:** All real-world use case examples presented in this article — including data fetching, processing, and modeling — are available at https://github.com/bokulich-publications/ritme_examples. The underlying datasets are publicly available from their original sources: NCBI SRA BioProject PRJEB5482^66^ (use case 1); the *Tara* Oceans companion resource^8^ at https://ocean-microbiome.embl.de/companion.html (use case 2); and https://github.com/SchlossLab/Sze_CRCMetaAnalysis_mBio_2018^75^ (use case 3).

## Acknowledgments

This work was supported by an ETH Zurich Doc.Mobility Fellowship awarded to Anja Adamov.

## Author Contributions

AA: Conceptualization, Data curation, Formal analysis, Funding acquisition, Investigation, Methodology, Software, Validation, Visualization, Writing – original draft, Writing – review & editing; CLM: Conceptualization, Methodology, Supervision, Writing – review & editing; NAB: Conceptualization, Methodology, Funding acquisition, Resources, Supervision, Writing – review & editing.

## Declaration of interests

The authors declare no competing interests.

## Declaration of generative AI and AI-assisted technologies in the writing process

During the preparation of this work, the authors used GitHub Copilot and Claude Code to assist with manuscript editing. After using these tools, the authors reviewed and edited the content as needed and take full responsibility for the content of the published article.

## Document S1. Supplemental information

**Table S1.** K-fold per-fold training style in *ritme* by trainable type.

|  | Non-iterative | Iterative |
| --- | --- | --- |
| Trainables | linreg, logreg, rf,<br>rf_class, trac | xgb, xgb_class,<br>nn_reg, nn_class, nn_corn |
| K-fold execution | Parallel | Sequential |

**Table S2.**
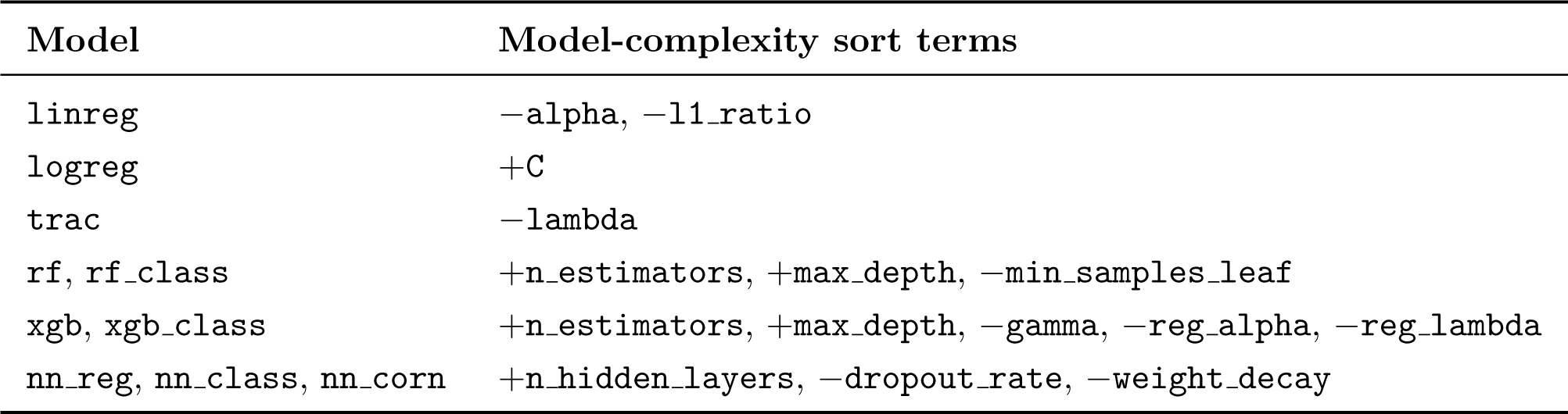
Definition of model complexity used by the one-standard-error rule for selecting parsimonious models. Terms are listed in decreasing priority order: each successive term resolves residual ties among configurations matched on all higher-priority terms. Each term +*h* (resp.*−h*) denotes that smaller (resp. larger) values of hyperparameter *h* correspond to simpler models. Model algorithm names and hyperparameters are defined in Tables 1 and S4.

**Figure S1.**
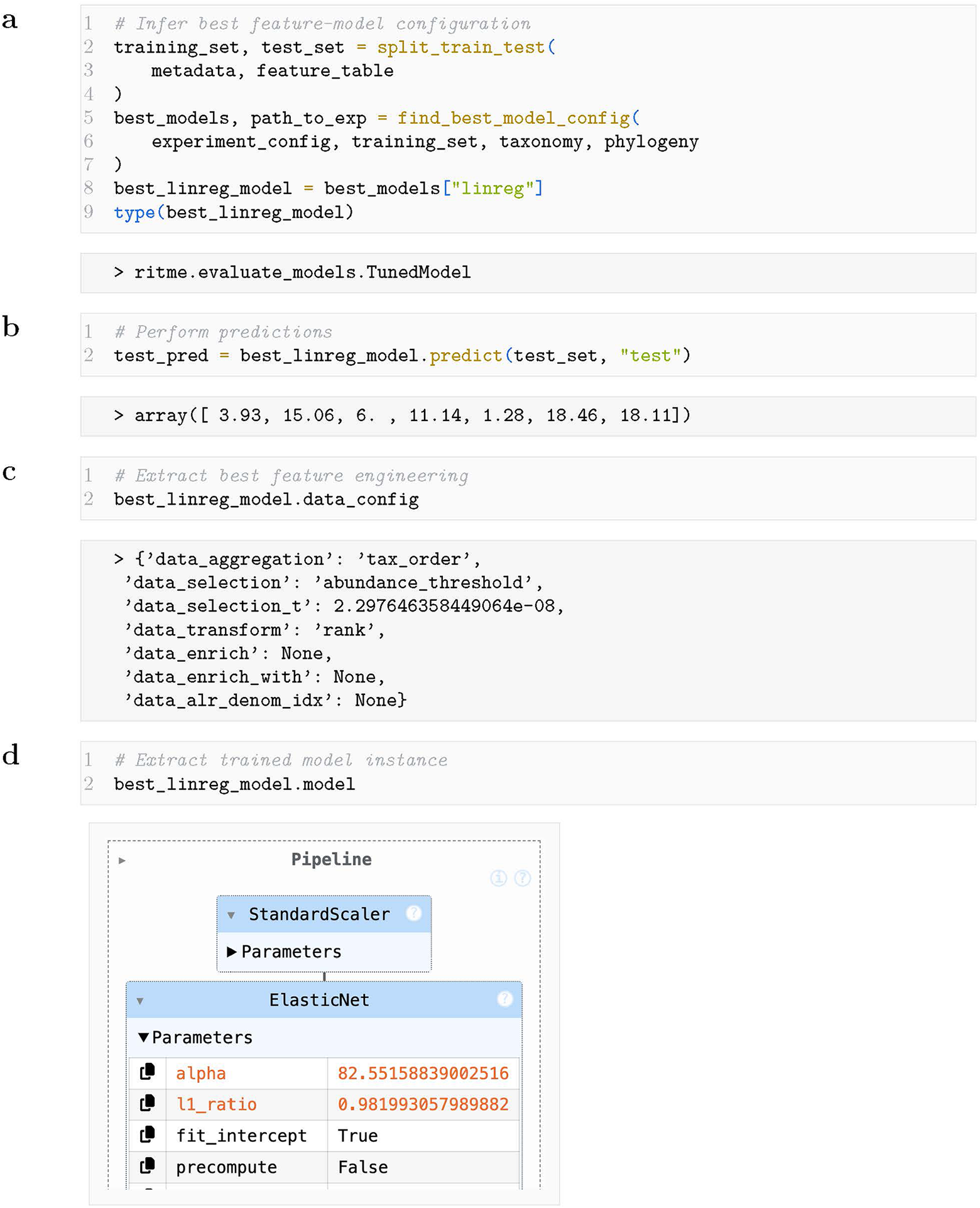
*ritme*’s Python API usage examples. (**a**) Finding the best feature-model configuration and the custom-made model object returned; (**b**) prediction on any dataset with the same feature structure; (**c**) extraction of the optimal feature engineering approach; and (**d**) the trained estimator.

**Figure S2.**
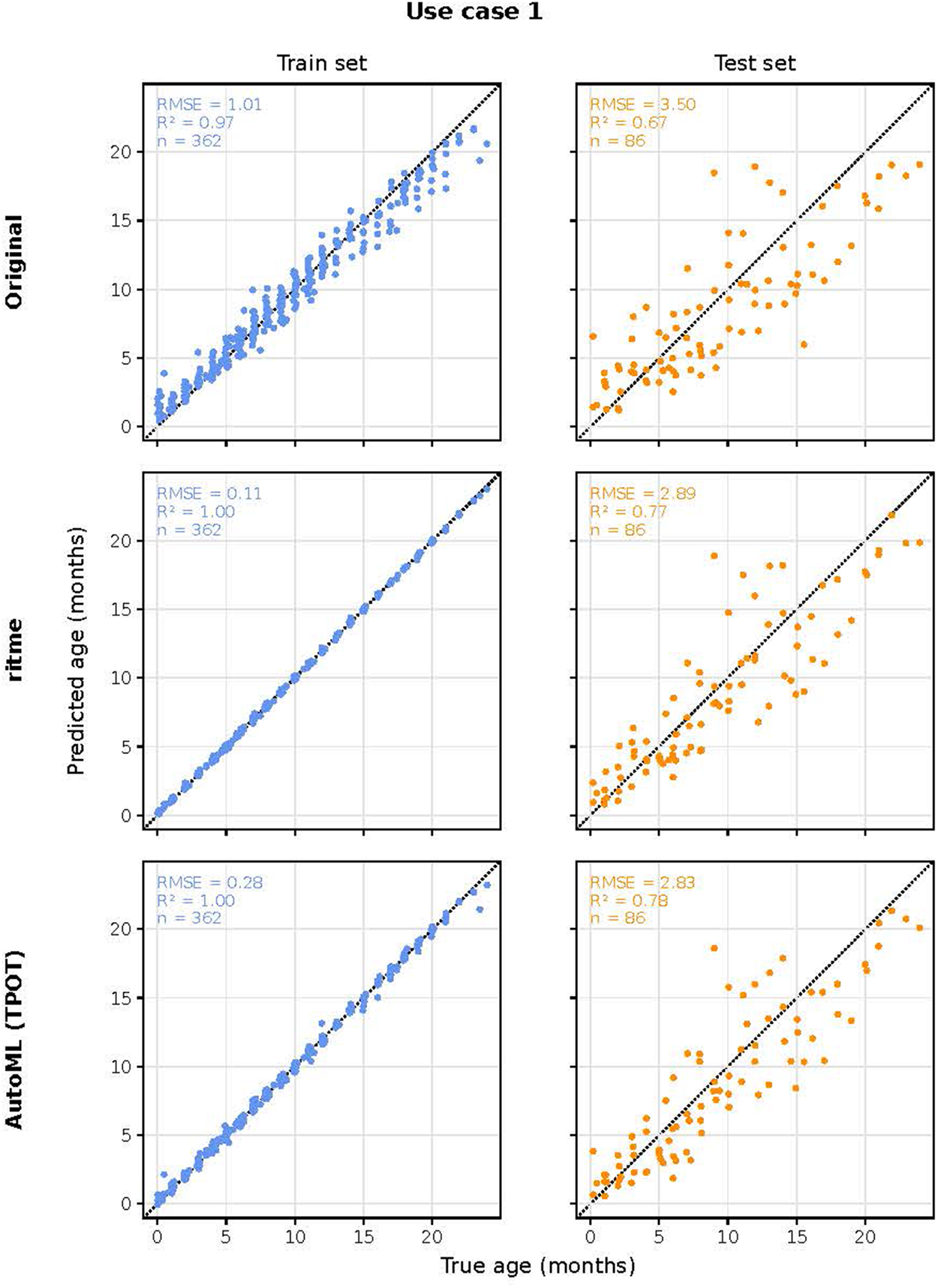
Train and held-out test set performance for use case 1. Shown are the original published approach^66^, *ritme*’s best feature-model configuration, and the strongest AutoML baseline *TPOT*, evaluated on the same train-test split of the infant age dataset (Table 2). Each panel plots predicted against true infant age with the identity line dashed; annotations report the split’s root mean squared error (RMSE), coefficient of determination (R^2^) and number of samples (n).

**Figure S3.**
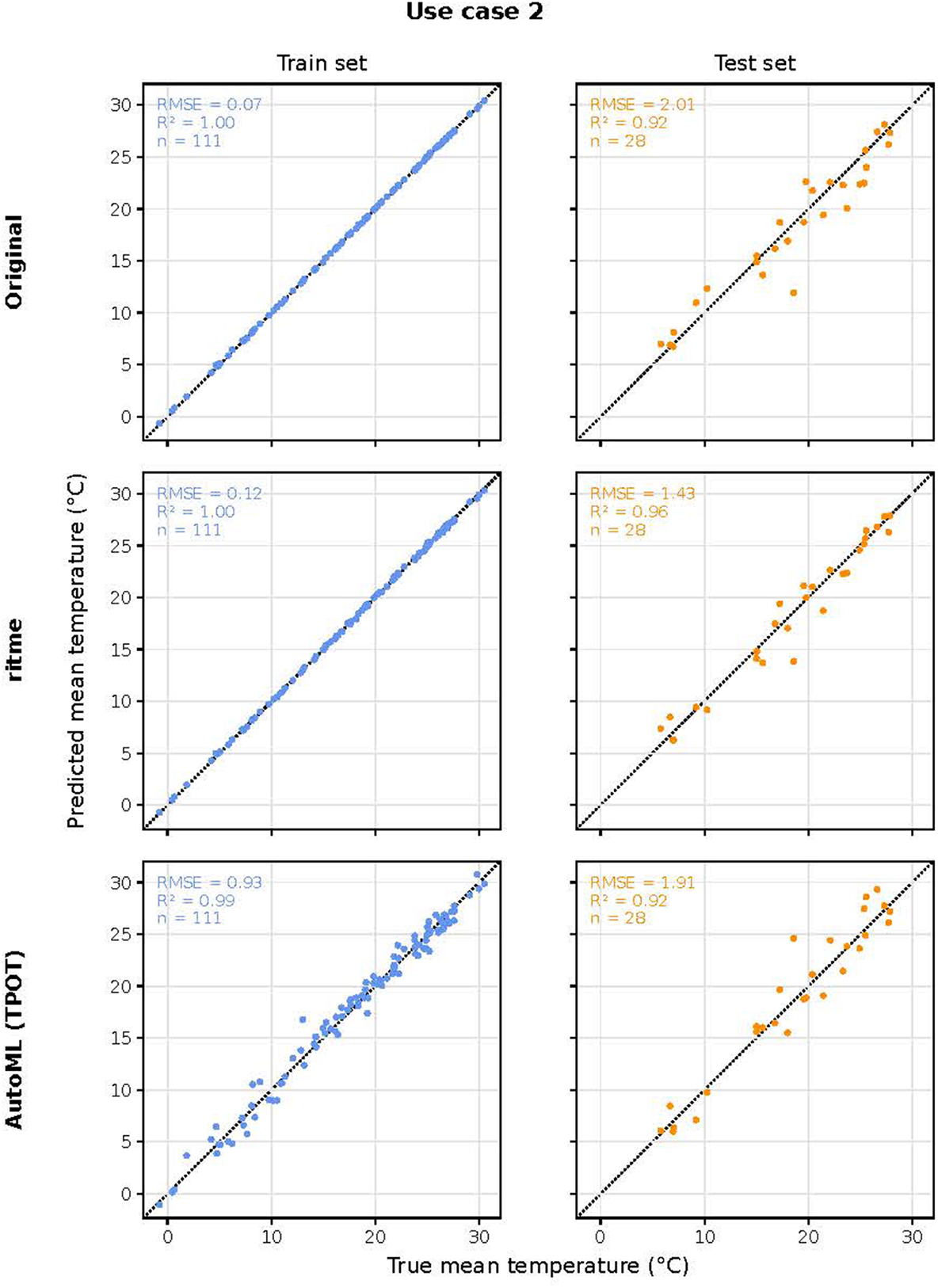
Train and held-out test set performance for use case 2. Shown are the original published approach^8^, *ritme*’s best feature-model configuration, and the strongest AutoML baseline *TPOT*, evaluated on the same train-test split of the ocean temperature dataset (Table 2). Each panel plots predicted against true ocean temperature with the identity line dashed; annotations report the split’s root mean squared error (RMSE), coefficient of determination (R^2^) and number of samples (n).

**Figure S4.**
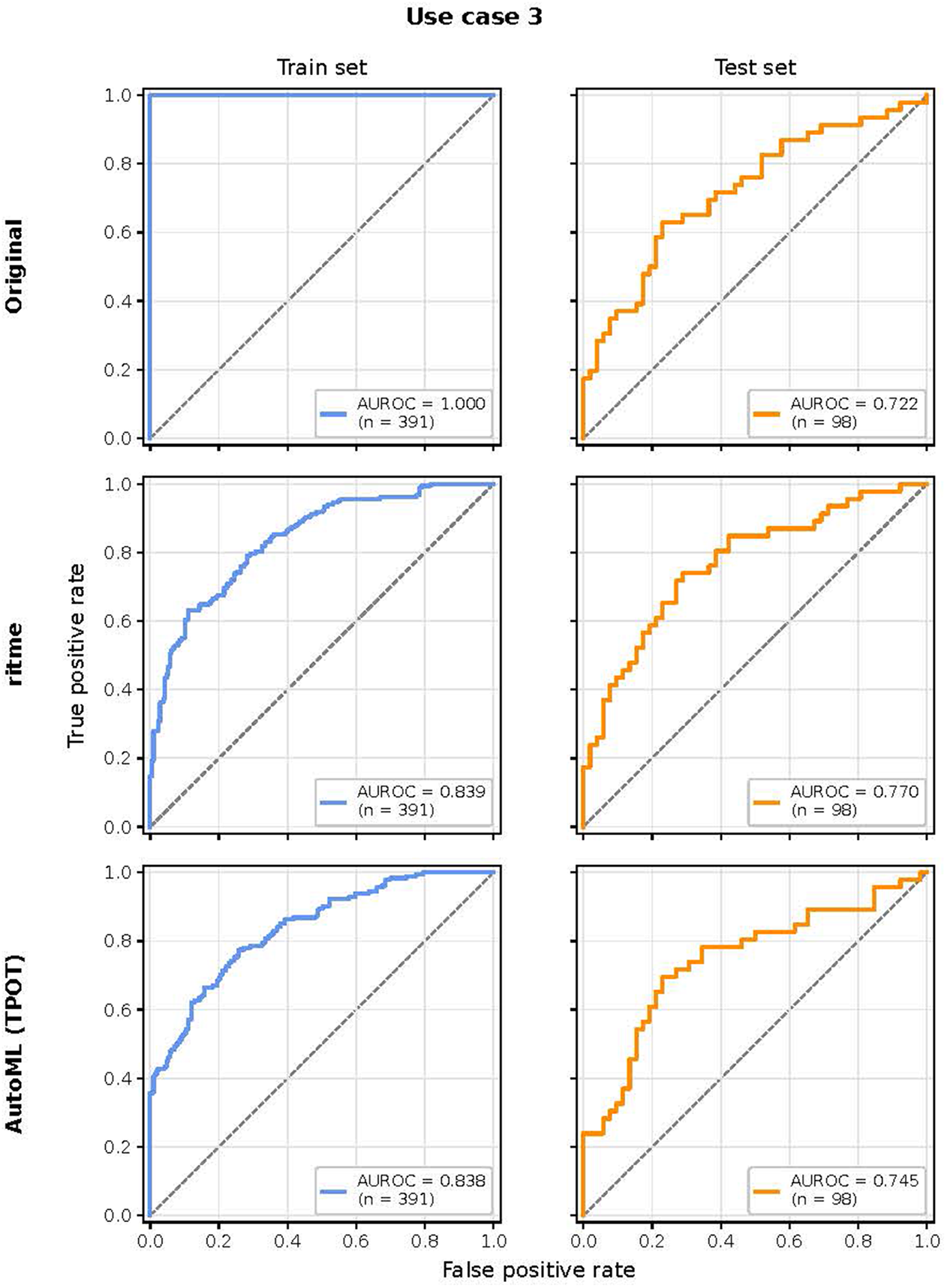
Train and held-out test set receiver operating characteristic curves for use case 3. Shown are the original published approach^75^, *ritme*’s best feature-model configuration, and the strongest AutoML baseline *TPOT*, evaluated on the same train-test split of the colorectal neoplasia dataset (Table 2). Annotations report the split’s area under the ROC curve (AUROC) and number of samples (n).

**Figure S5.**
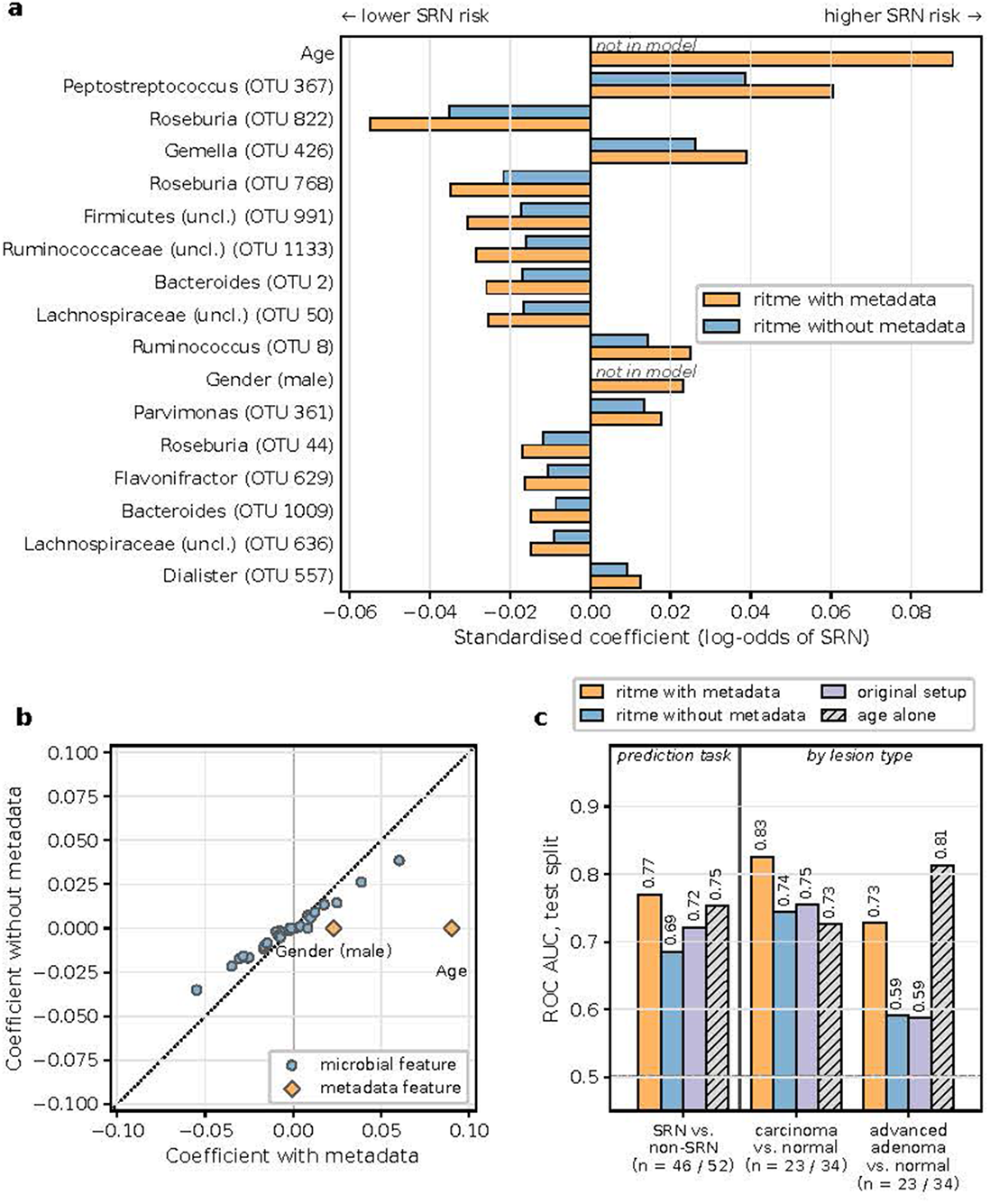
Comparison of the best *ritme* models with and without metadata enrichment for use case 3. Both models predict screen-relevant neoplasia (SRN). (**a**) Standardized logistic-regression coefficients of the features ranking in either model’s top 15 by absolute coefficient, ordered by the with-metadata model. (**b**) Coefficients of all features with a non-zero weight in at least one model. (**c**) Held-out test set ROC-AUC of the two *ritme* models, the original published set-up^75^, and age alone, for the SRN prediction task and for each SRN lesion type against normal colons. ROC-AUC = area under the receiver operating characteristic curve.

**Figure S6.**
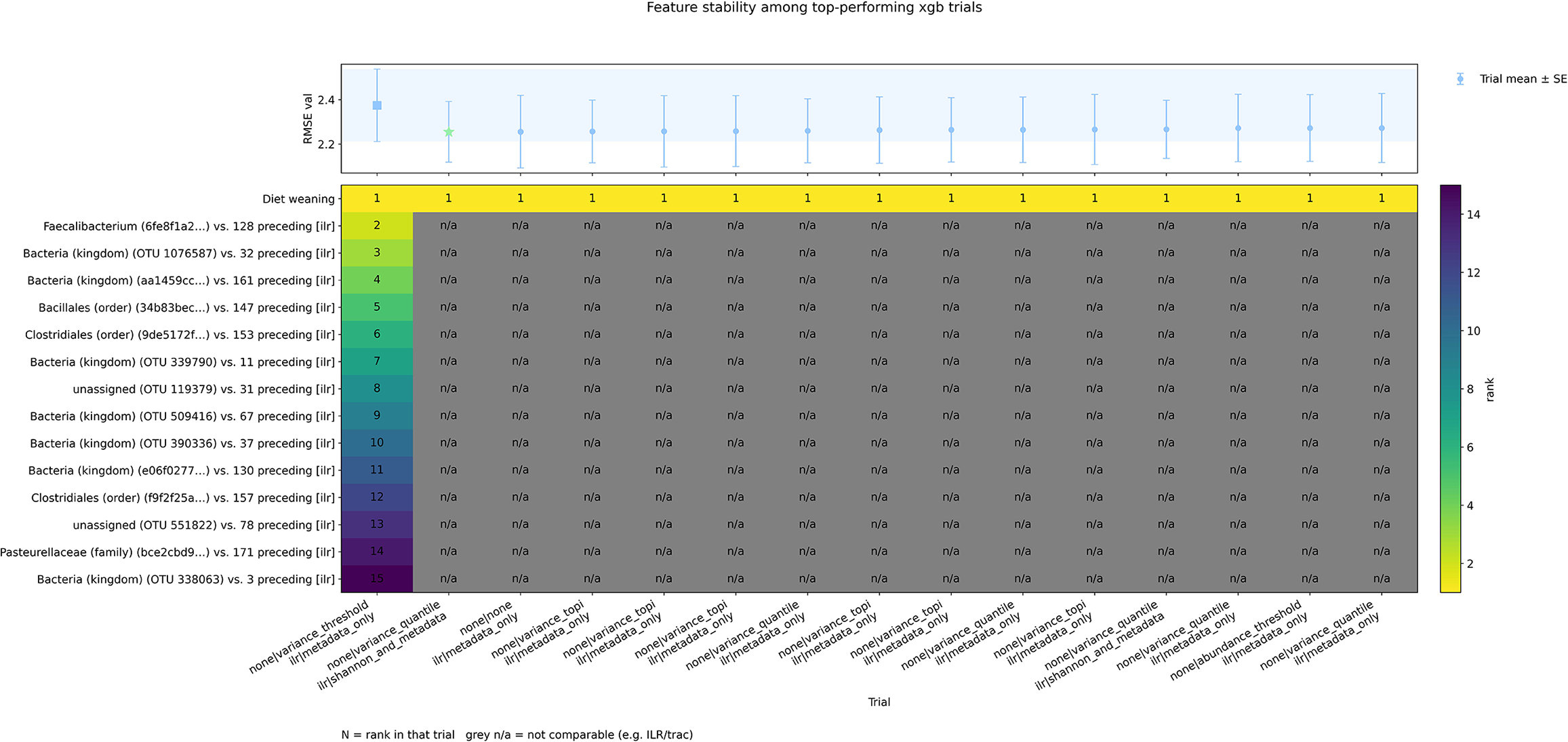
Feature-importance stability for use case 1 (xgb). Computed with explain stability. Top: mean cross-validation RMSE *±* standard error (SE) of the band of comparably performing trials, whose means lie within *±*1 SE of the reference trial, i.e. the selected best model (leftmost column, square marker; band capped at 15 trials: the reference plus the 14 next-best); the star marks the best mean and the shaded area spans the reference*±*1 SE. Columns are labeled by feature-engineering configuration (aggregation | selection | transform | enrichment). Bottom: rank of each of the reference’s top 15 features in every trial’s own importance ranking (1 = most important). All band trials use the isometric log ratio (ilr) transform, whose coordinates depend on each trial’s selected features; microbial features are therefore not comparable across trials (gray), and the sole comparable feature, the metadata covariate diet weaning, ranks first in all trials. RMSE = root mean squared error.

**Figure S7.**
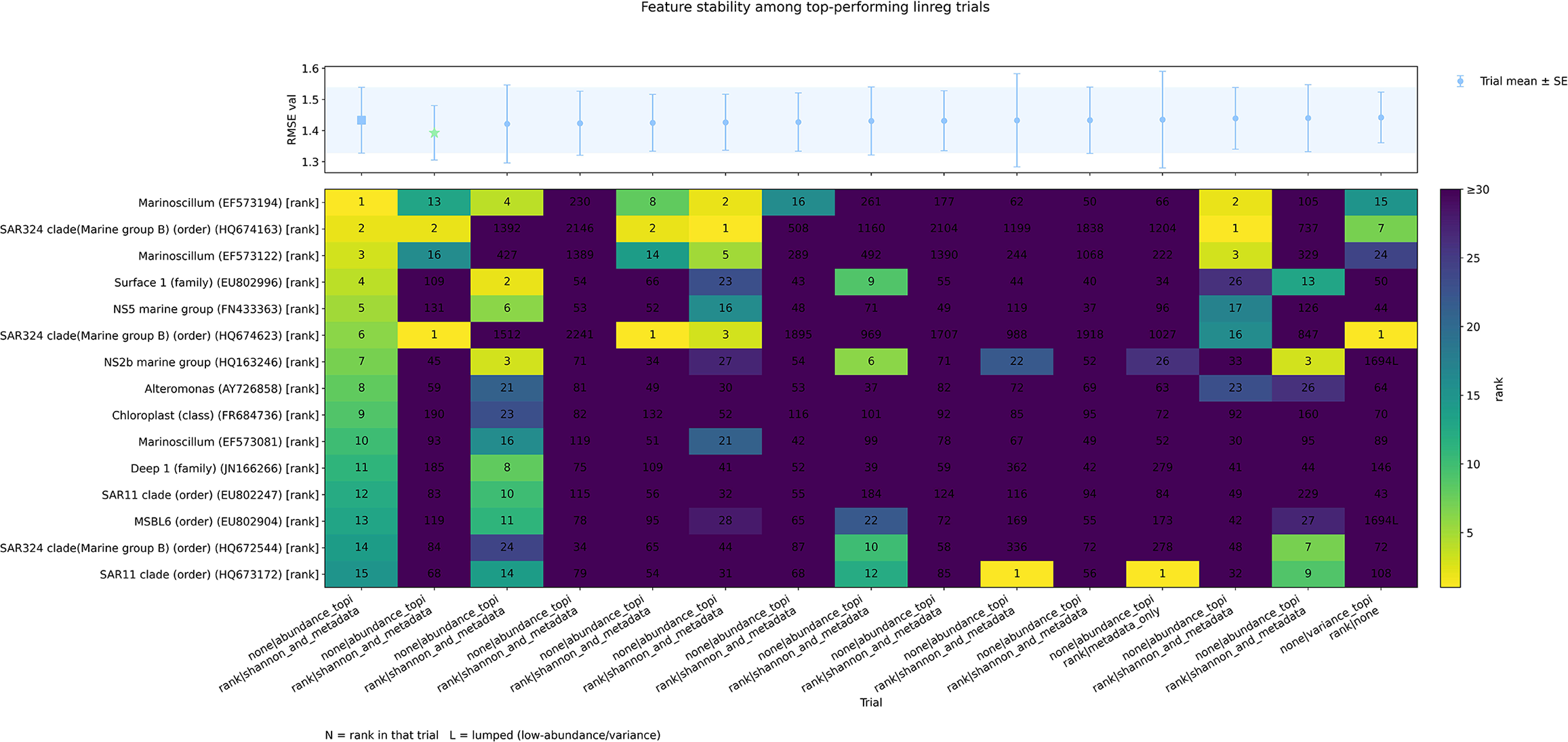
Feature-importance stability for use case 2 (linreg). Computed with explain stability. Top: mean cross-validation RMSE *±* standard error (SE) of the band of comparably performing trials, whose means lie within *±*1 SE of the reference trial, i.e. the selected best model (leftmost column, square marker; band capped at 15 trials: the reference plus the 14 next-best); the star marks the best mean and the shaded area spans the reference *±*1 SE. Columns are labeled by feature-engineering configuration (aggregation | selection | transform | enrichment). Bottom: rank of each of the reference’s top 15 features, matched across trials through their source features, in every trial’s own importance ranking (1 = most important). Cell annotations report each feature’s rank in the corresponding trial; ranks *≥* 30 share the darkest color, and the suffix L marks a feature matched into a trial’s lumped low-abundance or low-variance bucket. RMSE = root mean squared error.

**Figure S8.**
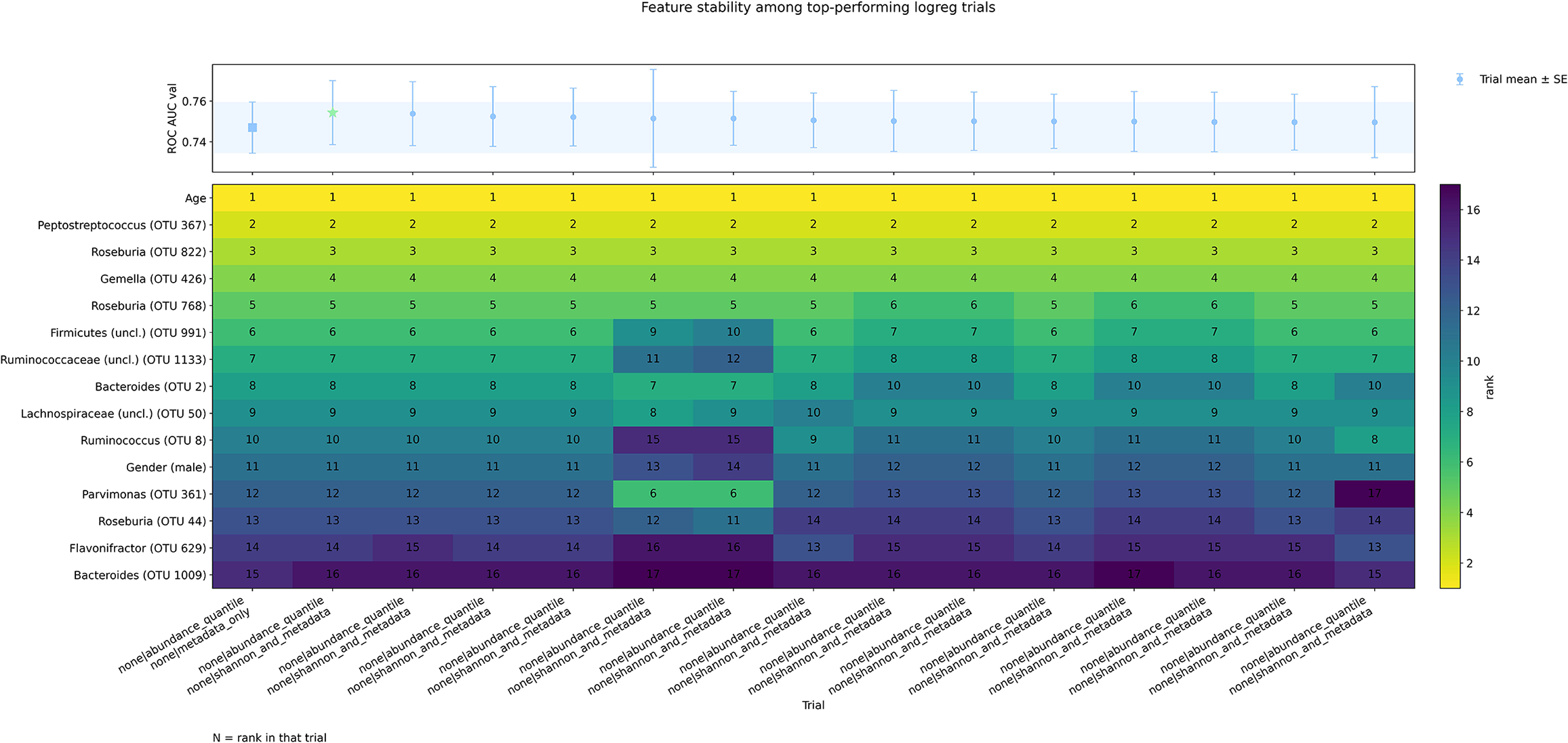
Feature-importance stability for use case 3 (logreg). Computed with explain stability. Top: mean cross-validation ROC-AUC *±* standard error (SE) of the band of comparably performing trials, whose means lie within *±*1 SE of the reference trial, i.e. the selected best model (leftmost column, square marker; band capped at 15 trials: the reference plus the 14 next-best); the star marks the best mean and the shaded area spans the reference *±*1 SE. Columns are labeled by feature-engineering configuration (aggregation | selection | transform | enrichment). Bottom: rank of each of the reference’s top 15 features in every trial’s own importance ranking (1 = most important). At least 13 of the reference’s top 15 features, led by the metadata covariate age, rank within each trial’s top 15. ROC-AUC = area under the receiver operating characteristic curve.

**Table S3.** Number and values of categorical (cat) and conditional continuous (c-cont.) hyperparameters for each feature engineering method in *ritme*. The conditional hyperparameter values, depicted in brackets, are considered only when their corresponding categorical parent parameter is selected.

| Name | # cat | # c-cont. | Values |
| --- | --- | --- | --- |
| Aggregation | 1 | 0 | None, tax_class, tax_order, tax_family, tax_genus |
| Selection | 1 | 1 | None |
|  |  |  | abundance_topi (*i*), variance_topi (*i*) |
|  |  |  | abundance_ith (*i*), variance_ith (*i*) |
|  |  |  | abundance_quantile (*q*), variance_quantile (*q*)<br>abundance_threshold (*t*), variance_threshold (*t*) |
| Transform | 1 | 0 | None, clr, ilr, alr, pa, rank |
| Enrichment | 1 | 0 | None, shannon, shannon_and_metadata, metadata_only |

**Figure S9.**
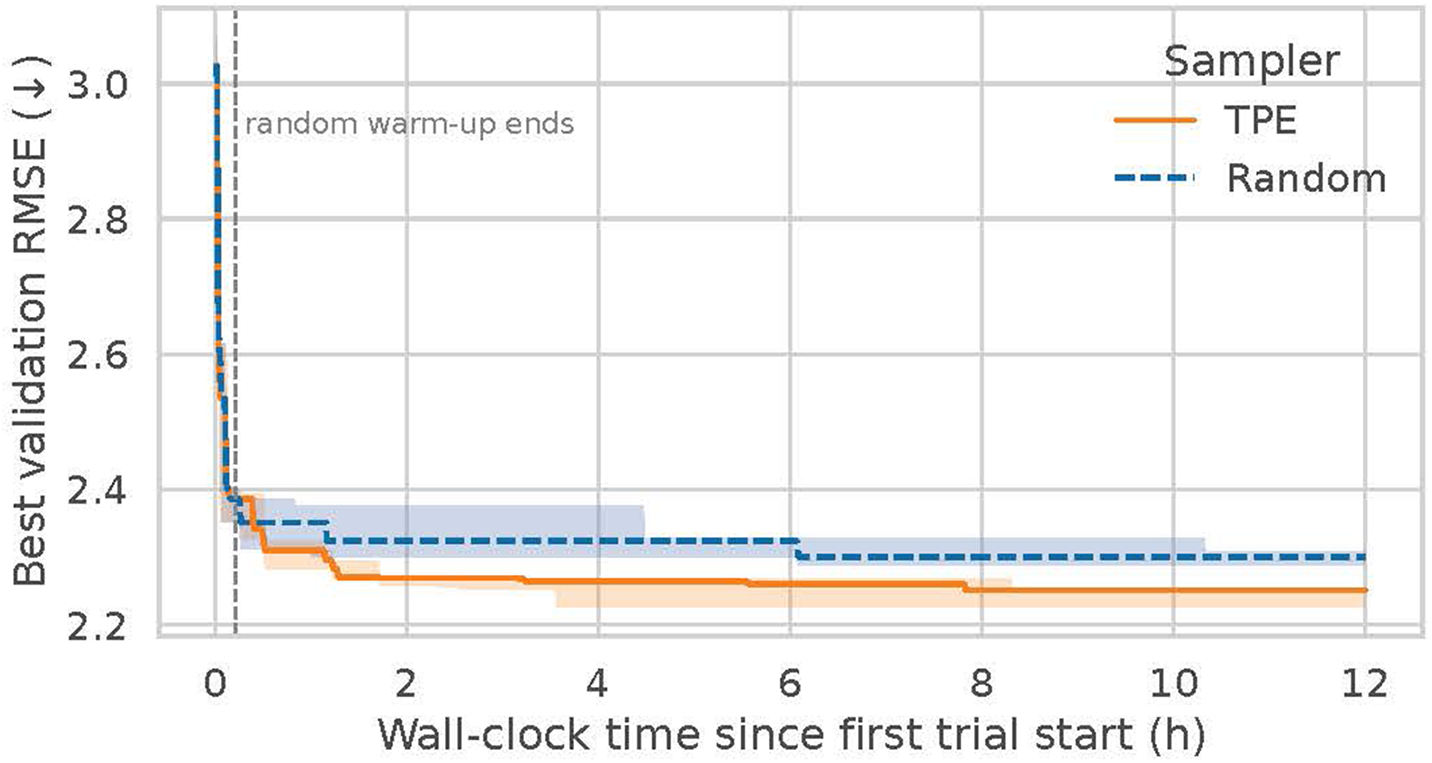
Search efficiency of *ritme*’s default TPE sampler for use case 1. Best validation RMSE achieved so far for the xgb model type (per-trial mean across K validation folds) against wall-clock time since the first trial start, compared to a purely random sampler. Each arm was run for 12 hours with three seeds; lines show the median across seeds and shaded bands the seed minimum–maximum. The dashed line marks the end of TPE’s adaptive random warm-up (75 trials; see “Search for best feature-model configuration”). RMSE = root mean squared error, TPE = Tree-structured Parzen Estimator.

**Table S4.** Number (#) and names of categorical (cat.) and continuous (cont.) hyperparameters for each predictive model in *ritme*. The neural network parameter n units hl*x* represents the number of neurons in the *x*-th hidden layer, where n hidden layers specifies the total number of hidden layers. All remaining parameter names match those of the original package implementations. Asterisk-marked parameters (*) were categorized to allow None as a hyperparameter option. Dagger-marked parameters (†) were discretized to reduce the hyperparameter search space. Evaluated default ranges for each hyperparameter are listed in Table S5.

| Model | # |  | Hyperparameter names |  |
| --- | --- | --- | --- | --- |
|  | cat | cont. | cat. | cont. |
| `linreg` | 0 | 2 | - | `alpha`, `l1_ratio` |
| `logreg` | 1 | 2 | `penalty` | `C`, `l1_ratio` |
| `rf`,<br>`rf_class` | 3 | 5 | `max_depth*`,<br>`max_features*`<br>`bootstrap` | `n_estimators`,<br>`min_samples_split`,<br>`min_samples_leaf`,<br>`min_weight_fraction_leaf`,<br>`min_impurity_decrease` |
| `trac` | 0 | 1 | - | `lambda` |
| `xgb`,<br>`xgb_class` | 0 | 10 | - | `n_estimators`, `max_depth`,<br>`min_child_weight`, `subsample`,<br>`eta`, `num_parallel_tree`,<br>`gamma`, `reg_alpha`,<br>`reg_lambda`, `colsample_bytree` |
| `nn_reg`,<br>`nn_class`,<br>`nn_corn` | `n_hidden_layers`<br>+ 3 | 5 | `n_units_hlx†`,<br>`learning_rate†`,<br>`batch_size†`,<br>`epochs†` | `n_hidden_layers`,<br>`dropout_rate`, `weight_decay`,<br>`early_stopping_patience`,<br>`early_stopping_min_delta` |

**Table S5.** Hyperparameters and their default evaluated ranges for each predictive model in *ritme*. Categorical hyperparameters list discrete evaluated values; continuous hyperparameters show the sampled interval [min, max] with log indicating log-uniform sampling and int indicating that only integer values are considered.

| Model | Categorical | Continuous |
| --- | --- | --- |
| linreg | – | alpha: $[10^{-5}, 10^2]$ (log)<br>l1_ratio: $[0, 1]$ |
| logreg | penalty: {"l1", "l2", "elasticnet"} | C: $[10^{-4}, 10^4]$ (log)<br>l1_ratio: $[0, 1]$ |
| rf,<br>rf_class | max_depth: {2, 4, 8, 16, 32, None}<br>max_features: {None, "sqrt", "log2"<br>0.1, 0.2, 0.3, 0.5}<br>bootstrap: {True, False} | n_estimators: $[20, 3000]$ (int)<br>min_samples_split: $[10^{-3}, 10^{-1}]$ (log)<br>min_samples_leaf: $[10^{-3}, 10^{-1}]$ (log)<br>min_weight_fraction_leaf: $[0, 10^{-2}]$<br>min_impurity_decrease: $[0, 0.5]$ |
| trac | – | lambda: $[10^{-3}, 1]$ (log) |
| xgb,<br>xgb_class | – | n_estimators: $[20, 3000]$ (int)<br>max_depth: $[2, 10]$ (int)<br>min_child_weight: $[0, 4]$ (int)<br>subsample: $[0.7, 1.0]$<br>eta: $[10^{-2}, 0.3]$ (log)<br>num_parallel_tree: $[1, 4]$ (int)<br>gamma: $[10^{-3}, 5]$ (log)<br>reg_alpha: $[10^{-10}, 1]$ (log)<br>reg_lambda: $[10^{-10}, 1]$ (log)<br>colsample_bytree: $[0.3, 1.0]$ |
| nn_reg,<br>nn_class,<br>nn_corn | n_units_hlx: {32, 64, 128, 256, 512}<br>learning_rate: {1e-2, 5e-3, 1e-3<br>5e-4, 1e-4, 5e-5, 1e-5}<br>batch_size: {32, 64, 128, 256}<br>epochs: {10, 50, 100, 200} | n_hidden_layers: $[1, 30]$ (int)<br>dropout_rate: $[0, 0.5]$<br>weight_decay: $[10^{-6}, 10^{-2}]$ (log)<br>early_stopping_patience: $[2, 30]$ (int)<br>early_stopping_min_delta: $[10^{-5}, 10^{-2}]$ (log) |

**Table S6.**
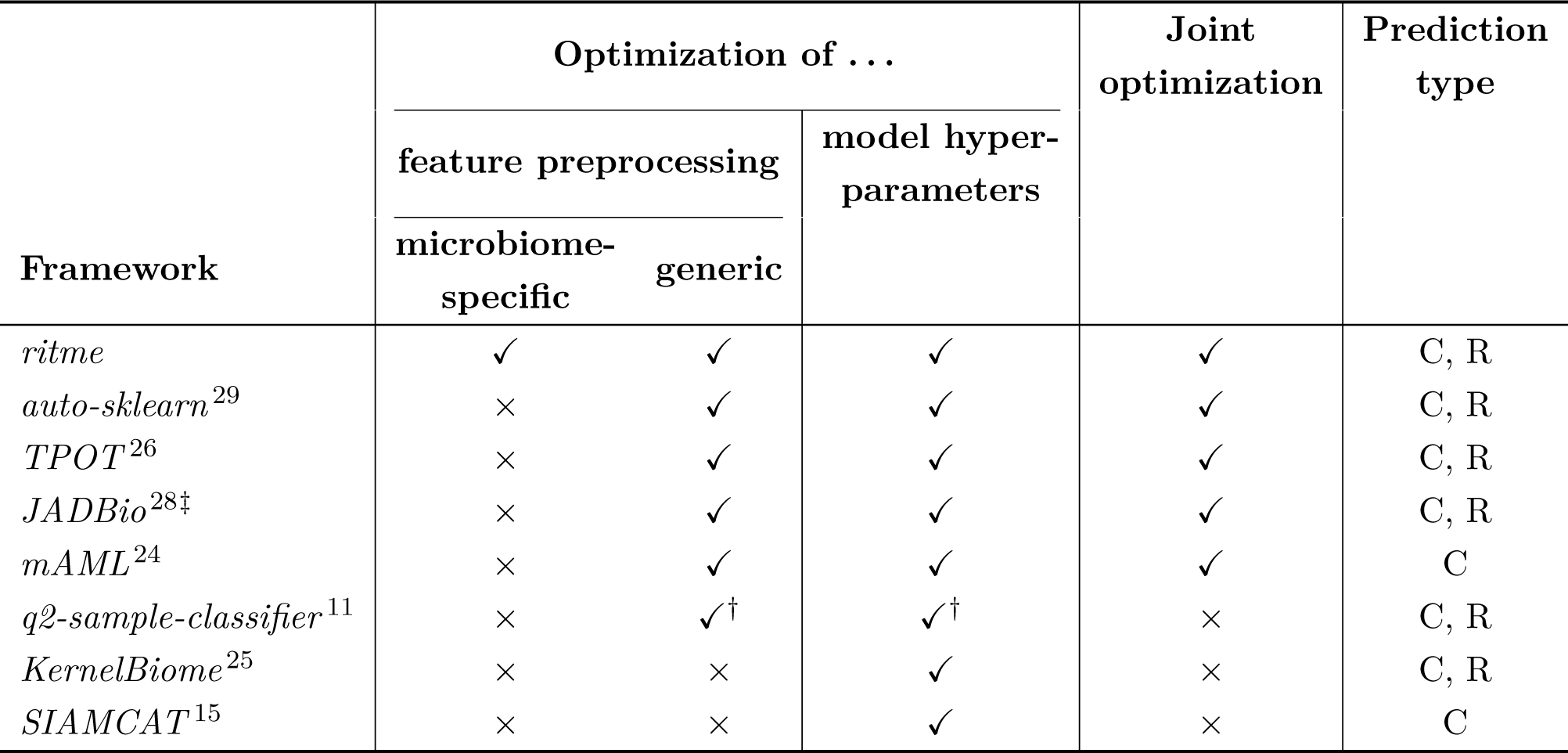
Frameworks applicable to supervised prediction from microbial feature count tables, characterized by the pipeline components they optimize. Frameworks that search feature preprocessing jointly with model class and hyperparameters (column “Joint optimization”) qualified as AutoML baselines for comparison with *ritme*; of these, *JADBio* (‡) was not retained owing to its proprietary distribution. ✓ = optimized; *×* = not optimized, although fixed, user-selected functionality may be available; † = disabled by default. C = classification, R = regression.

**Figure S10.**
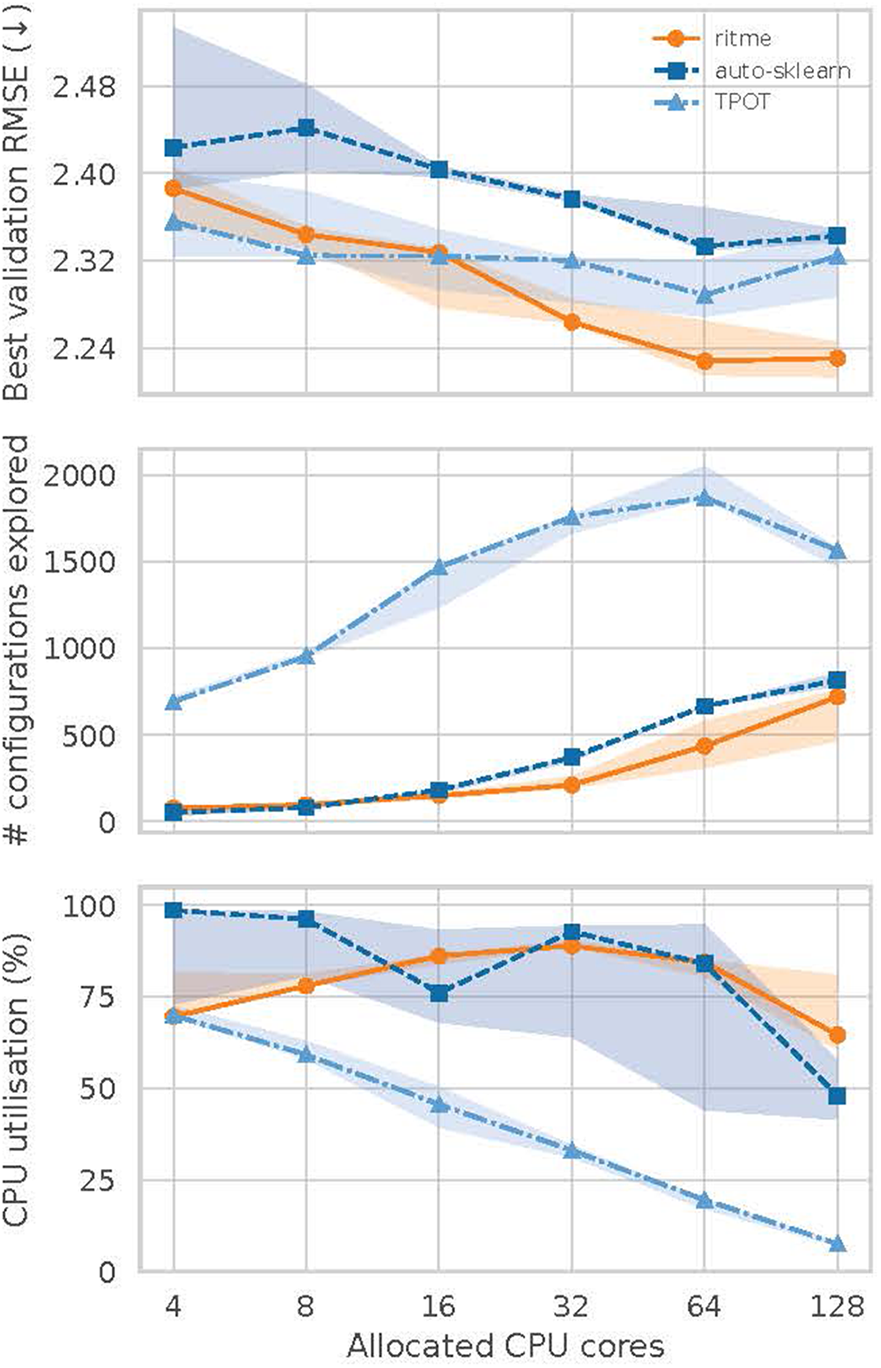
Compute scaling of *ritme* and the generic AutoML baselines for use case 1. Best validation RMSE (top), number of explored configurations (middle), and CPU utilization derived from the cluster’s accounting records (bottom) of *ritme*, *auto-sklearn*, and *TPOT* as a function of allocated CPU cores. Per allocation, each arm was run for a fixed two-hour search budget with three seeds on the same node type; all arms received the identical train/test split and metadata enrichment, were evaluated with the same grouped 5-fold cross-validation, and the baselines were restricted to the estimator family matching *ritme*’s winning model type (see Methods). Lines show the median across seeds and shaded bands the seed minimum–maximum. RMSE = root mean squared error, TPE = Tree-structured Parzen Estimator.

## Notes

https://github.com/adamovanja/ritme

### Competing Interest Statement

The authors have declared no competing interest.

### Summary of Updates

additional benchmarks & new feature-stability method in ritme

